# DIABLO: from multi-omics assays to biomarker discovery, an integrative approach

**DOI:** 10.1101/067611

**Authors:** Amrit Singh, Casey P. Shannon, Benoît Gautier, Florian Rohart, Michaël Vacher, Scott J. Tebbutt, Kim-Anh Lê Cao

## Abstract

Systems biology approaches, leveraging multi-omics measurements, are needed to capture the complexity of biological networks while identifying the key molecular drivers of disease mechanisms. We present DIABLO, a novel integrative method to identify multi-omics biomarker panels that can discriminate between multiple phenotypic groups. In the multi-omics analyses of simulated and real-world datasets, DIABLO resulted in superior biological enrichment compared to other integrative methods, and achieved comparable predictive performance with existing multi-step classification schemes. DIABLO is a versatile approach that will benefit a diverse range of research areas, where multiple high dimensional datasets are available for the same set of specimens. DIABLO is implemented along with tools for model selection, and validation, as well as graphical outputs to assist in the interpretation of these integrative analyses (http://mixomics.org/).

## Background

Technological improvements have allowed for the collection of data from different molecular compartments (*e.g.*, gene expression, methylation status, protein abundance) resulting in multiple omics (multi-omics) data from the same set of biospecimens (*eg.*, transcriptomics, proteomics, metabolomics). The large number of omic variables compared to the limited number of available biological samples presents a computational challenge when identifying the key drivers of disease. Further, technological limitations differ with respect to different omic platforms (*e.g.*, sequencing *vs.* mass spectrometry), and biological effect sizes differ with respect to different omic variable-types (e.g., methylation status *vs.* protein expression). Effective integrative strategies are needed, to extract common biological information spanning multiple molecular compartments that explains phenotypic variation. Already, systems biology approaches which incorporated data from multiple biological compartments, have shown improved biological insights compared to traditional single omics analyses [1–3]. This may be because single omics analyses cannot account for the interactions between omic layers and, consequently, are unable to reconstruct accurate molecular networks. These molecular networks are dynamic, changing under perturbed conditions such as disease, response to therapy, and environmental exposures. Therefore, adopting a holistic approach by integrating multi-omics data may bridge this information gap, and uncover networks that are representative of the underlying molecular mechanisms [4, 5].

Preliminary approaches to data integration included multi-step approaches that leveraged existing single-omics methods: multi-omics data were concatenated, or ensembles of single omics models created [6]. These approaches can be biased towards certain omics data types, however, and do not account for interactions between omic layers [7, 8]. Recently, more sophisticated integrative approaches have been proposed (**Supplementary Fig. 1**) [4,9–12]. They can be broadly divided into unsupervised analyses, which identify coherent relationships across multi-omics datasets when samples are unlabeled, and supervised analyses, which identify multi-omics patterns that discriminate between known phenotypic sample groups. However these supervised strategies are unable to capture the shared information across multiple biological domains when identifying the key molecular drivers associated with a phenotype. Such methods are needed to capture the dynamic nature of molecular networks under various disease conditions and ultimately provide robust biomarkers that are both biologically and clinically relevant.

To address these knowledge gaps, we introduce DIABLO, a method that incorporates information across high dimensional multi-omics data while discriminating phenotypic groups. DIABLO uncovers robust biomarkers of dysregulated disease processes that span multiple functional layers. We demonstrate the capabilities and versatility of DIABLO both in simulated and real-world data, integrating multi-omics datasets to identify relevant biomarkers of various diseases. DIABLO is available through the mixOmics data integration toolkit (www.mixomics.org [12]) which contains a wide range of multivariate methods for the exploration and integration of high dimensional biological datasets.

## Results

DIABLO (**<u>D</u>**ata **<u>I</u>**ntegration **<u>A</u>**nalysis for **<u>B</u>**iomarker discovery using **<u>L</u>**atent c**<u>O</u>**mponents) maximizes the common or correlated information between multiple omics (multi-omics) datasets while identifying the key omics variables (mRNA, miRNA, CpGs, proteins, metabolites, *etc.*) and characterizing the disease sub-groups or phenotypes of interest. DIABLO uses Projection to Latent Structure models (PLS) [13], and extends both sparse PLS-Discriminant Analysis [14] to multi-omics analyses and sparse Generalized Canonical Correlation Analysis [15] to a supervised analysis framework. In contrast to existing penalized matrix decomposition methods [16], DIABLO is a component-based method (or a dimension reduction technique) that transforms each omic dataset into latent components and maximizes the sum of pairwise correlations between latent components (user-defined) and a phenotype of interest [17]. DIABLO is, therefore, an integrative classification method that builds predictive multi-omics models that can be applied to multi-omics data from new samples to determine their phenotype. Users can specify the number of variables to select from each dataset and visualize the omics data and the multi-omics panel into a reduced data. The method is highly flexible in the type of experimental design it can handle, ranging from classical single time point to cross-over and repeated measures studies. Modular-based analysis can also be incorporated using pathway-based module matrices [18] instead of the original omics matrices, as illustrated in one of our case studies.

### DIABLO selects correlated and discriminatory variables

Briefly, three omic datasets consisting of 200 samples (split equally over two groups) and 260 variables were generated by modifying the degree of correlation and discrimination, resulting in four types of variables: 30 correlated-discriminatory (corDis) variables, 30 uncorrelated-discriminatory (unCorDis) variables, 100 correlated-nondiscriminatory (corNonDis) variables, and 100 uncorrelated-nondiscriminatory (unCorNonDis) variables (**Supplementary Note, Supplementary Fig. 2**). Three integrative classification methods were applied to generate multi-omic biomarkers panels of 90 variables each (30 variables from each omic dataset): a DIABLO model with either a full design (where the correlation between all pairwise combinations of datasets, as well as between each dataset and the phenotypic outcome, were maximised) or the null design (where only the correlation between each dataset and the phenotypic outcome was maximised, **see Methods**), a concatenation-based sPLSDA classifier which consists of naively combining all datasets into one, and an ensemble of sPLSDA classifiers where a separate sPLSDA classifier was fitted for each omics dataset and the consensus predictions were combined using a majority vote scheme (see **Supplementary Fig. 3**). The purpose of the simulation study was to compare DIABLO models with existing multi-step integrative classifiers with respect to the error rate and types of variables selected as part of the multi-omic biomarker panels. A secondary aim was to determine the effect of design matrix on the resulting multi-omic biomarker panels identified using DIABLO.

The concatenation, ensemble and DIABLO_null classifiers performed similarly across the various noise and fold-change thresholds. At lower noise levels (simulated using a multivariate normal distribution with mean of zero and standard deviation of 0.2 or 0.5) the DIABLO_full classifier had a slightly higher error rate compared to the other approaches (Fig. 1a), but consistently selected mostly correlated and discriminatory (corDis) variables, unlike the other integrative classifiers (Fig. 1b). All methods behaved similarly with respect to the error rate and types of variables selected at higher noise thresholds (simulated using a multivariate normal distribution with mean of zero and standard deviation of 1 or 2). This simulation highlights how the design (connection between datasets) affects the flexibility of the DIABLO model, resulting in a trade-off between discrimination or correlation. DIABLO_null focused on selecting discriminatory variables and disregarded most of the correlation between datasets (null design), whereas DIABLO_full selected highly correlated variables across all datasets. Since the variables selected by DIABLO_full reflect the correlation structure between biological compartments, we hypothesized that they might provide a balance between prediction accuracy and biological insight.

**Figure 1.**
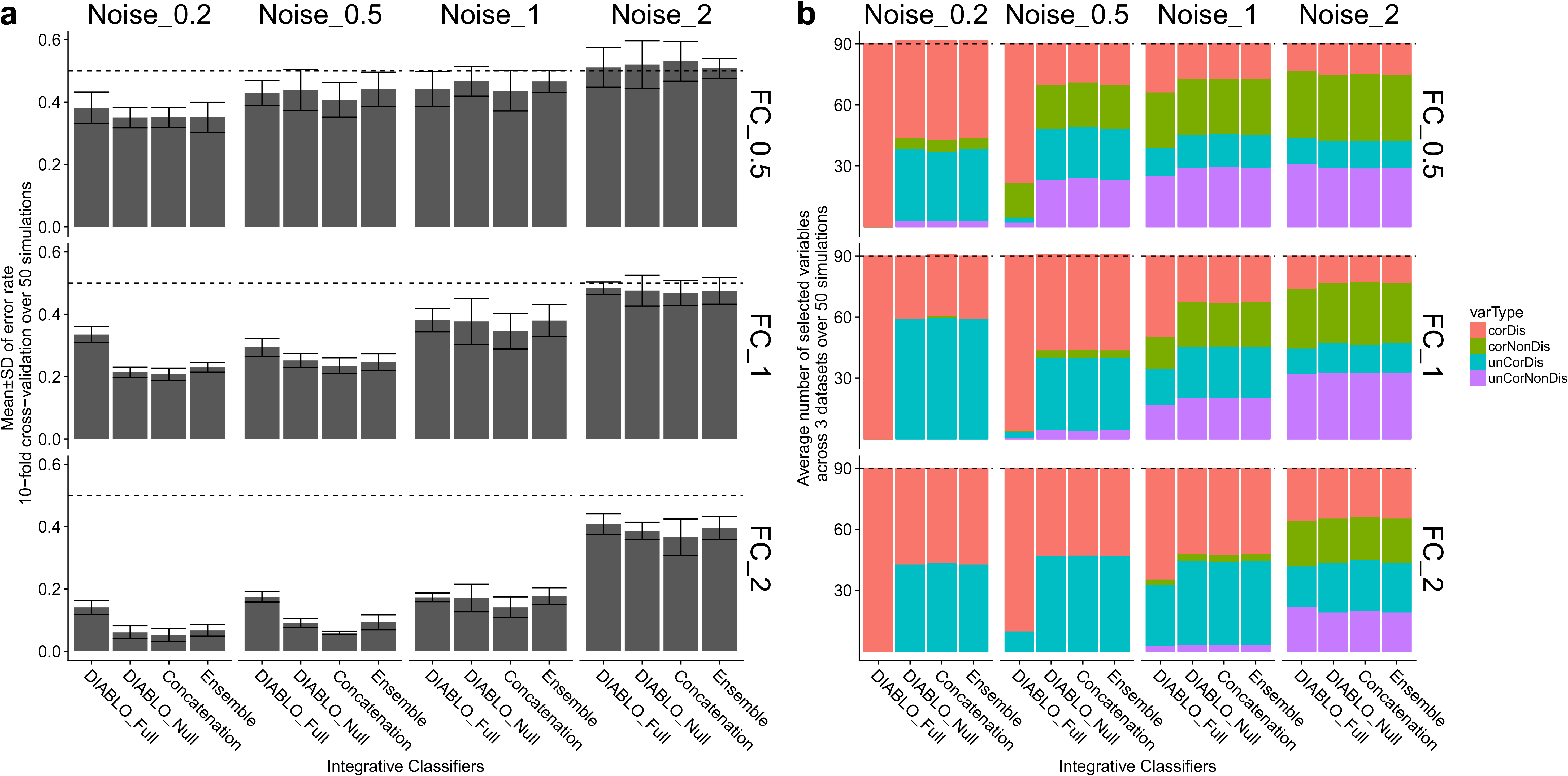
Simulation study: performance assessment and benchmarking. Simulated datasets included different types of variables: correlated & discriminatory (corDis); uncorrelated & discriminatory (unCorDis); correlated & nondiscriminatory (corNonDis) and uncorrelated & nondiscriminatory (unCorNonDis) for different fold-changes between sample groups and different noise levels (**see Supplementary Note**). Integrative classifiers included DIABLO with either the full or null design, concatenation and ensemble-based sPLSDA classifiers and were all trained to select 90 variables across three multi-omics datasets. **a)** Classification error rates (10-fold cross-validation averaged over 50 simulations). Dashed line indicates a random performance (error rate = 50%). All methods perform similarly with the exception of DIABLO_full which has a higher error rate. **b)** Number of variables selected according to their type. DIABLO_full selected mainly variables that were correlated & discriminatory (corDis, red), whereas the other methods selected an equal number of correlated or uncorrelated discriminatory variables (corDis and unCorDis, red and blue).

### DIABLO identifies molecular networks with superior biological enrichment

To assess this, we turn to real biological datasets. We applied various integrative approaches to cancer multi-omics datasets (mRNA, miRNA, and CpG) – colon, kidney, glioblastoma (gbm) and lung – and identified multi-omics biomarker panels that were predictive of high and low survival times (Table 1). We then compared the network properties and biological enrichment of the selected features across approaches.

**Table 1.** Overview of multi-omics datasets analyzed for method benchmarking and in two case studies. The breast cancer case study includes training and test datasets for all omics types except proteins.

| Analysis | Dataset | Number of samples | Sample size in each subtype |  |  | Omics | Number of variables |
| --- | --- | --- | --- | --- | --- | --- | --- |
| **Benchmark cancer datasets (Wang et al. [3])** | Colon | 92 | High (33)<br>Low (59) |  |  | mRNA | 17,814 |
|  |  |  |  |  |  | miRNA | 312 |
|  |  |  |  |  |  | CpGs | 23,088 |
|  | Kidney | 122 | High (61)<br>Low (61) |  |  | mRNA | 17,665 |
|  |  |  |  |  |  | miRNA | 329 |
|  |  |  |  |  |  | CpGs | 24,960 |
|  | Glioblastoma (gbm) | 213 | High (105)<br>Low (108) |  |  | mRNA | 12,042 |
|  |  |  |  |  |  | miRNA | 534 |
|  |  |  |  |  |  | CpGs | 1,305 |
|  | Lung | 106 | High (53)<br>Low (53) |  |  | mRNA | 12,042 |
|  |  |  |  |  |  | miRNA | 353 |
|  |  |  |  |  |  | CpGs | 23,074 |
| **Case study 1 (The Cancer Genome Atlas ) [19]** | Breast cancer | 989 |  | Train | Test | mRNA | 16,851 |
|  |  |  | Basal | 76 | 102 | miRNA | 349 |
|  |  |  | Her2 | 38 | 40 | CpGs | 9,482 |
|  |  |  | LumA | 188 | 346 | Proteins | Train: 115<br>Test: 0 |
|  |  |  | LumB | 77 | 122 |  |  |
| **Case study 2 (Singh et al. [20,21])** | Asthma | 28 | Pre (14)<br>Post (14) |  |  | Cell-types | 9 |
|  |  |  |  |  |  | mRNA-modules | 229 |
|  |  |  |  |  |  | metabolite-modules | 60 |

Multi-omics biomarker panels were developed using component-based integrative approaches that also performed variable selection: supervised methods included concatenation and ensemble schemes using the sPLSDA classifier [14], and DIABLO with either the null or full design (DIABLO_null, and DIABLO_full); unsupervised approaches included sparse generalized canonical correlation analysis [15] (sGCCA), Multi-Omics Factor Analysis (MOFA), and Joint and Individual Variation Explained (JIVE) [23] (see **Supplementary Note** for parameter settings). Both supervised and unsupervised approaches were considered in order to compare and contrast the types of omics-variables selected, network properties and biological enrichment results. A distinction was made between DIABLO models in which the correlation between omics datasets was not maximized (DIABLO_null) and those when the correlation between omics datasets was maximized (DIABLO_full).

Each multi-omics biomarker panel included 180 features (60 features of each omics type across 2 components). Approaches generally identified distinct sets of features. Fig. 2a depicts the distinct and shared features between the seven multi-omics panels obtained from the unsupervised (purple, sGCCA, MOFA and JIVE) and supervised (green, Concatenation, Ensemble, DIABLO_null and DIABLO_full) methods. Supervised methods selected many of the same features (blue), but DIABLO_full had greater feature overlap with unsupervised methods (orange). The level of connectivity of each of the seven multi-omics panels was assessed by generating networks from the feature adjacency matrix at various Pearson correlation coefficient cut-offs (Fig. 2b). At all cut-offs, unsupervised approaches produced networks with greater connectivity (number of edges) compared to supervised approaches. In addition, biomarker panels identified by DIABLO_full, were more similar to those identified by unsupervised approaches, including high graph density, low number of communities and large number of triads, indicating that DIABLO_full identified discriminative sets of features that were tightly correlated across biological compartments (**Supplementary Fig. 4**). For example, Fig. 2c (upper panel) depicts the networks of all multi-omics biomarker panels for the colon cancer dataset, which show higher modularity (a limited number of large clusters of variables; circled) for the DIABLO_full and the unsupervised approaches as compared to the supervised ones. The corresponding component plots show a clear separation between the high and low survival groups for the panels derived using supervised approaches, whereas the unsupervised approaches could not segregate the survival groups [Fig. 2c (lower panel), see **Supplementary Fig. 5 and 6** for other cancer datasets].

**Figure 2.**
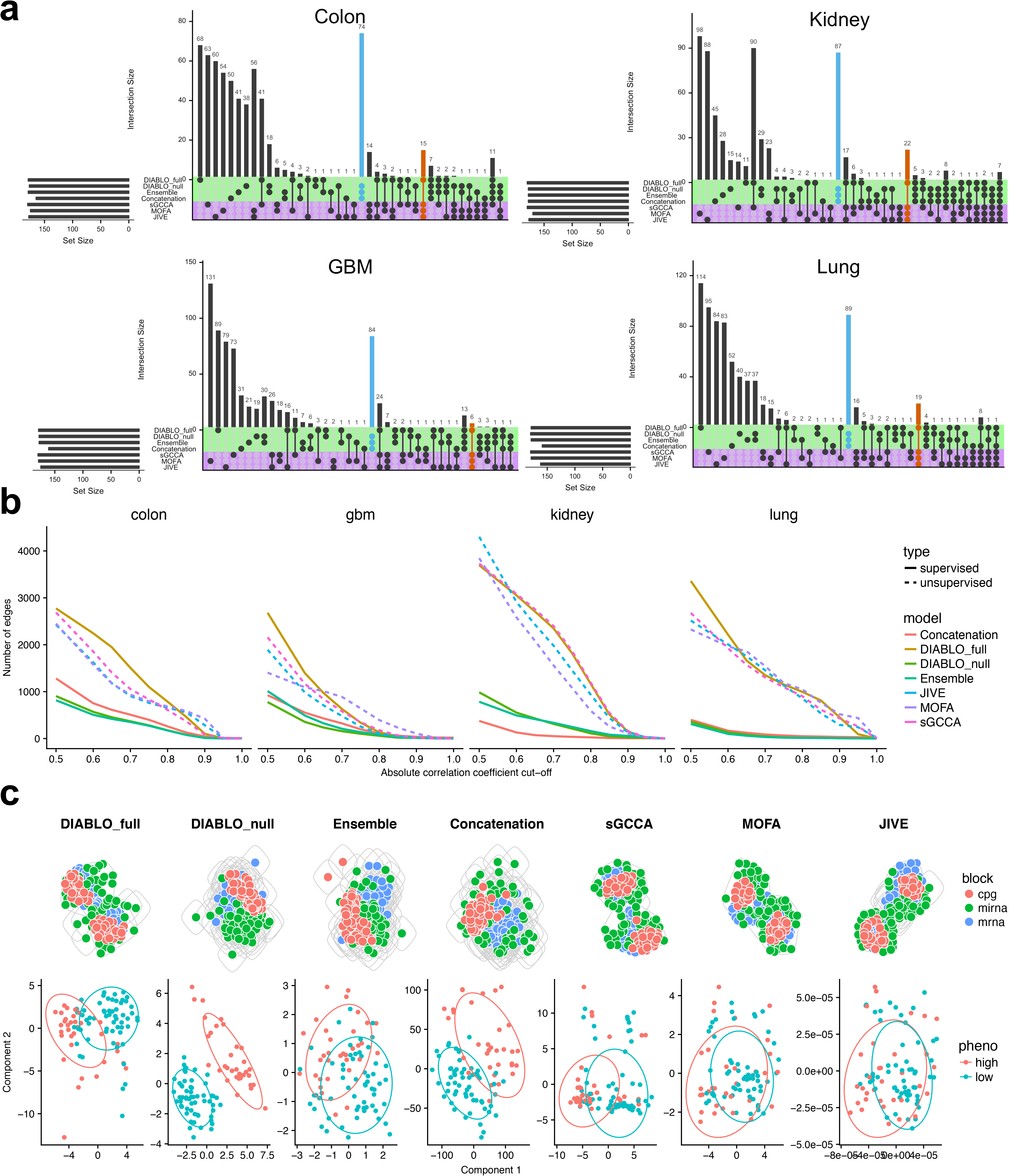
Benchmarking integrative methods using multi-omics biomarker panels for different cancers. **a)** Overlap of selected features using both supervised (green) and unsupervised approaches (purple): a strong overlap was observed between the supervised approaches with the exception of DIABLO_full (blue bars) which showed more similarity to unsupervised methods (dark orange bars). **b)** Number of edges within each panel network at various Pearson correlation cut-offs: unsupervised approaches panels were more connected than those from supervised approaches, with the exception of DIABLO_full which led to a highly-connected panel. An edge is present if the association between two omic variables is greater than a given correlation cut-off. **c)** Upper panel: network modularity of each multi-omic biomarker panel for colon cancer showed that unsupervised approaches and DIABLO_full resulted in a few groups of highly connected features, whereas supervised approaches identified networks with many groups of sparsely connected features. Lower panel: component plots depicting the clear separation of subjects in the high and low survival groups for supervised methods as opposed to the unsupervised methods.

Finally, we carried out gene set enrichment analysis on each multi-omics biomarker panel (using gene symbols of mRNAs and CpGs) against 10 gene set collections (see **Methods**) and tabulated the number of significant (FDR=5%) gene sets (Table 2). The DIABLO_full model identified the greatest number of significant gene sets across the 10 gene set collections and generally ranked higher than the other methods in the colon (7 collections), gbm (5 collections) and lung (5 collections) cancer datasets, whereas JIVE outperformed all other methods in the kidney cancer datasets (6 collections). Unlike all other approaches considered, DIABLO_full, which aimed to explain both the correlation structure between multiple omics layers and a phenotype of interest, implicated the greatest number of known biological gene sets (pathways/functions/processes *etc.*).

**Table 2.** Number of significant gene sets for each integrative method and benchmarking cancer dataset. Best performing method is indicated in the shaded cell. Each row represents a gene set collection (**see Methods** for details, FDR = 5%).

|  |  | Unsupervised, integrative |  |  | Supervised, non-integrative |  |  | Supervised, integrative |
| --- | --- | --- | --- | --- | --- | --- | --- | --- |
| disease | collection | JIVE | MOFA | sGCCA | Concatenation | Ensemble | DIABLO_null | DIABLO_full |
| Colon | BTM | 0 | 4 | 0 | 0 | 0 | 0 | 23 |
|  | C1 | 0 | 0 | 0 | 0 | 0 | 0 | 0 |
|  | C2 | 15 | 14 | 5 | 12 | 3 | 21 | 113 |
|  | C3 | 8 | 5 | 14 | 11 | 2 | 6 | 0 |
|  | **C4** | 0 | 1 | 0 | 1 | 2 | 1 | 46 |
|  | **C5** | 19 | 36 | 147 | 7 | 0 | 0 | 216 |
|  | **C6** | 0 | 0 | 0 | 0 | 0 | 0 | 0 |
|  | **C7** | 1 | 87 | 11 | 61 | 10 | 62 | 218 |
|  | **H** | 0 | 0 | 0 | 0 | 0 | 2 | 7 |
|  | **TISSUES** | 2 | 12 | 0 | 0 | 0 | 0 | 16 |
|  | **TOTAL** | 45 | 159 | 177 | 92 | 17 | 92 | 639 |
| **Gbm** | **BTM** | 0 | 0 | 19 | 10 | 9 | 10 | 30 |
|  | **C1** | 0 | 0 | 0 | 0 | 0 | 0 | 0 |
|  | **C2** | 275 | 337 | 193 | 258 | 358 | 312 | 426 |
|  | **C3** | 94 | 64 | 37 | 14 | 15 | 15 | 34 |
|  | **C4** | 49 | 43 | 68 | 47 | 50 | 62 | 125 |
|  | **C5** | 825 | 708 | 706 | 526 | 669 | 776 | 693 |
|  | **C6** | 22 | 25 | 18 | 30 | 24 | 24 | 21 |
|  | **C7** | 460 | 82 | 526 | 432 | 173 | 147 | 869 |
|  | **H** | 12 | 8 | 8 | 19 | 23 | 20 | 19 |
|  | **TISSUES** | 18 | 29 | 21 | 10 | 12 | 14 | 44 |
|  | **TOTAL** | 1755 | 1296 | 1596 | 1346 | 1333 | 1380 | 2261 |
| **Kidney** | **BTM** | 1 | 0 | 0 | 0 | 0 | 0 | 0 |
|  | **C1** | 0 | 0 | 1 | 0 | 0 | 0 | 1 |
|  | **C2** | 42 | 33 | 7 | 10 | 5 | 15 | 4 |
|  | **C3** | 8 | 80 | 1 | 4 | 35 | 23 | 1 |
|  | **C4** | 17 | 6 | 0 | 7 | 1 | 3 | 0 |
|  | **C5** | 157 | 110 | 1 | 55 | 27 | 46 | 0 |
|  | **C6** | 0 | 0 | 0 | 0 | 0 | 0 | 0 |
|  | **C7** | 0 | 74 | 15 | 93 | 13 | 10 | 18 |
|  | **H** | 6 | 3 | 0 | 1 | 0 | 1 | 0 |
|  | **TISSUE S** | 2 | 0 | 0 | 0 | 0 | 0 | 0 |
|  | **TOTAL** | 233 | 306 | 25 | 170 | 81 | 98 | 24 |
| **Lung** | **BTM** | 0 | 0 | 0 | 0 | 0 | 2 | 0 |
|  | **C1** | 0 | 0 | 0 | 1 | 0 | 0 | 1 |
|  | **C2** | 4 | 17 | 2 | 0 | 0 | 1 | 33 |
|  | **C3** | 48 | 20 | 57 | 50 | 26 | 21 | 19 |
|  | **C4** | 17 | 0 | 47 | 0 | 0 | 18 | 13 |
|  | **C5** | 35 | 127 | 42 | 0 | 25 | 22 | 193 |
|  | **C6** | 1 | 0 | 1 | 3 | 2 | 5 | 7 |
|  | **C7** | 18 | 13 | 78 | 0 | 7 | 72 | 100 |
|  | **H** | 0 | 2 | 0 | 0 | 1 | 0 | 0 |
|  | **TISSUE**<br>**S** | 0 | 0 | 0 | 0 | 0 | 9 | 20 |
|  | **TOTAL** | 123 | 179 | 227 | 54 | 61 | 150 | 386 |

### Case study 1: DIABLO identified known and novel multi-omics biomarkers of breast cancer subtypes

We next demonstrate that DIABLO can identify novel biomarkers in addition to biomarkers with known biological associations using a case study of human breast cancer. We applied our biomarker analysis workflow to breast cancer datasets to characterize and predict PAM50 breast cancer subtypes (**Supplementary Fig. 7**). After preprocessing and normalization of each omics data-type, the samples were divided into training and test sets (**Methods,** Table 1). The training data consisted of four omics-datasets (mRNA, miRNA, CpGs and proteins) whereas the test data included all remaining samples for which the protein expression data were missing. The optimal multi-omics biomarker panel size was identified using a grid approach where, for any given combination of variables, we assessed the classification performance using a 5-fold cross-validation repeated 5 times (**Supplementary Fig. 8**). The number of variables that resulted in the minimum balanced error rate were retained as previously described in [12]. The optimal multi-omics panel consisted of 45 mRNA, 45 miRNAs, 25 CpGs and 55 proteins selected across three components with a balanced error rate of 17.9±1.9%. This panel identified many variables with previously known associations with breast cancer, as assessed by looking at the overlap between the panel features and gene sets related to breast cancer based on the Molecular Signature database (MolSigDB) [23], miRCancer [24], Online Mendelian Inheritance in Man (OMIM) [25], and DriverDBv2 [26]. Figure 3a depicts the variable contributions of each omics-type indicated by their loading weight (variable importance). Variables not found in any database may represent novel biomarkers of breast cancer. Figure 3b shows the consensus and individual omics component plots based on this biomarker panel, along with 95% confidence ellipses obtained from the training data and superimposed with the samples from the test data. The majority of the samples were within the ellipses, suggesting a reproducible multi-omics biomarker panel from the training to the test set, that was predictive of breast cancer subtypes (balanced error rate = 22.9%). The consensus plot corresponded strongly with the mRNA component plot, depicting a strong separation of the Basal (error rate = 4.9%) and Her2 (error rate = 20%) subtypes. We observed a weak separation of Luminal A (LumA, error rate = 13.3%) and Luminal B (LumB, error rate = 53.3%) subtypes. Similarly, the heatmap showing the scaled expression of all features of the multi-omics biomarker panel, depicted a strong clustering of the Basal and Her2 samples whereas the Luminal A and B were mixed (Fig. 3c). Overall, the features of the multi-omics biomarker panel formed a densely connected network comprising of four communities where variables in each community (cluster) were densely connected with themselves and sparsely connected with other clusters (Fig. 3d). The largest cluster in Fig. 3d consisted of 72 variables; 20 mRNAs, 21 miRNAs, 15 CpGs and 16 proteins (red bubble) and was further investigated using gene set enrichment analysis. We identified many cancer-associated pathways (*e.g.* FOXM1 pathway, p53 signaling pathway), DNA damage and repair pathways (*e.g.* E2F mediated regulation of DNA replication, G2M DNA damage checkpoint) and various cell-cycle pathways (*e.g.* G1S transition, mitotic G1/G1S phases), demonstrating the ability of DIABLO to identify a biologically plausible multi-omics biomarker panel. This panel generalized to new breast cancer samples and implicated previously unknown molecular features in breast cancer, which could be further validated in experimental studies.

**Figure 3.**
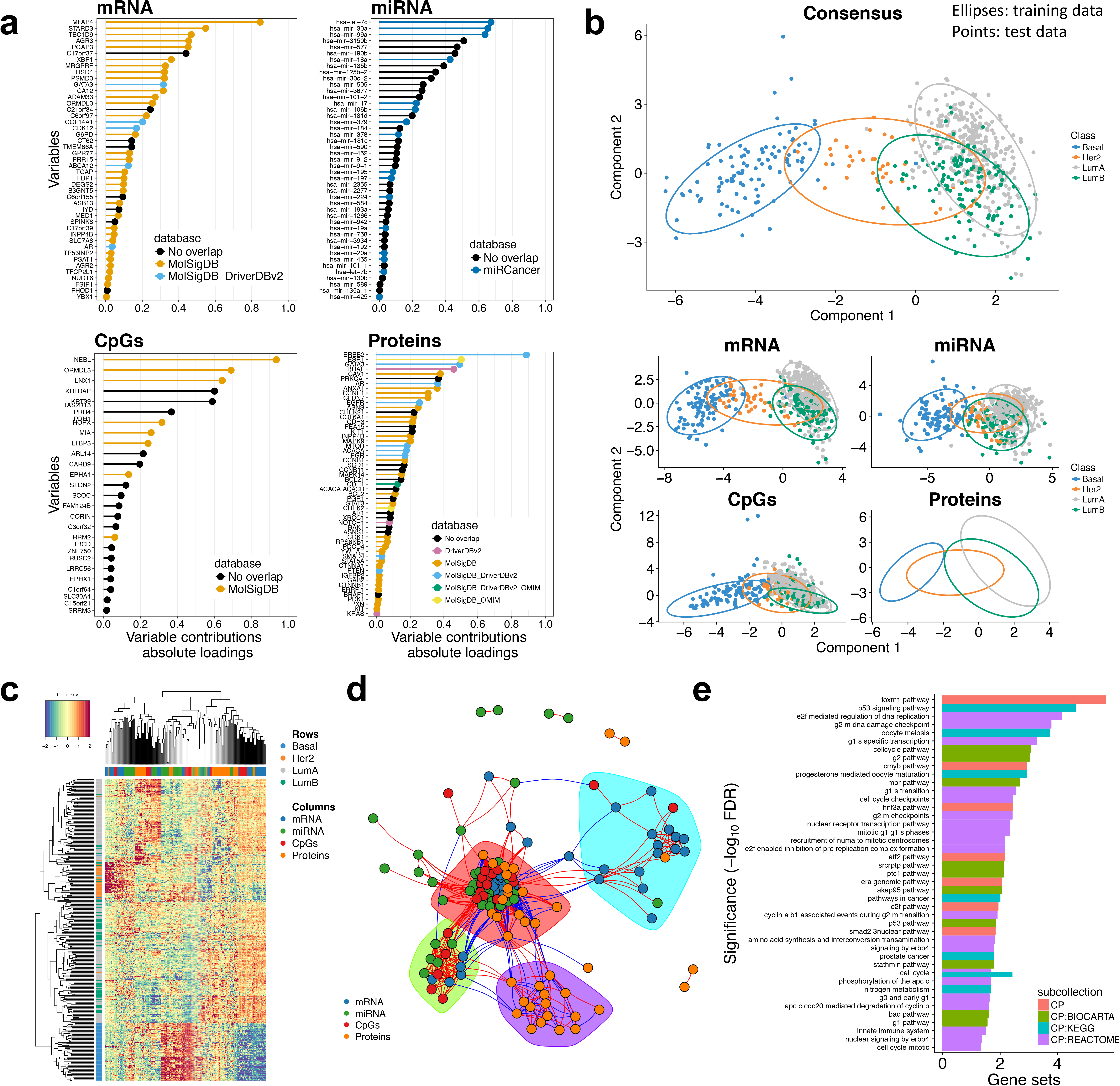
Identification of a multi-omics biomarker panel predictive of breast cancer subtypes. **a)** Variable contributions of each omics-type biomarker that are important to discriminate breast cancer subtypes. **b)** DIABLO component plots and the derived biomarker panel: 95% confidence ellipses were calculated from the training data set and points depict samples from the test set. **c)** Heatmap of the scaled expression of variable from the biomarker panel. **d)** Network visualization of the biomarker panel highlights correlated variables (Pearson correlation > |0.4|) and four communities based on edge betweeness scores. **e)** A gene set enrichment analysis was conducted on the largest community from d (red cluster) where many cancer related pathways were identified.

### Case study 2: DIABLO for repeated measures designs and module-based analyses

Next, we demonstrate the flexibility of DIABLO by extending its use to a repeated measures cross-over study [27], as well as incorporating module-based analyses that incorporate prior biological knowledge [28–30]. We use a small multi-omics asthma dataset, including pre and post intervention timepoints, to compare a DIABLO model that can account for repeated measures (multilevel DIABLO) with the standard DIABLO model as described above [20, 21]. An allergen inhalation challenge was performed as we previously described in [20, 21] in 14 subjects and blood samples were collected before (pre) and two hours after (post) challenge; cell-type frequencies, leukocyte gene transcript expression and plasma metabolite abundances were determined for all samples (Table 1). We observed a net decline in lung function after allergen inhalation challenge (**Supplementary Fig. 9**), and the goal of this study was to identify perturbed molecular mechanisms in the blood in response to allergen inhalation challenge. A module based approach (also known as eigengene summarization [18], **see Methods**) was used to transform both the gene expression and metabolite datasets into pathway datasets.

Consequently, each variable in those two datasets now represented the scaled pathway activity expression level for each sample instead of direct gene/metabolite expression. The mRNA dataset was transformed into a dataset of metabolic pathways (based on the Kyoto Encyclopedia of Genes and Genomes, KEGG) whereas the metabolite dataset was transformed into a metabolite pathway dataset based on annotations provided by Metabolon Inc. (Durham, North Carolina, USA) (Fig. 4a). To account for the repeated measures experimental design, a multilevel approach [27] was first used to isolate the within-sample variation from each dataset (see **Methods**), and then DIABLO was applied to identify a multi-omics biomarker panel consisting of cells, gene and metabolite modules that discriminated pre-from post-challenge samples. We contrast the resulting ‘multilevel DIABLO’ (mDIABLO) with a standard DIABLO model that disregards the paired nature of this study by comparing their cross-validation classification performances (Fig. 4b). mDIABLO outperformed DIABLO (AUC=98.5% vs. AUC=62.2%, leave-one-out cross-validation, **see Methods**), and we observed a greater degree of separation between the pre- and post-challenge samples for mDIABLO compared to DIABLO (Fig. 4c). Common features (pathways) were identified across omics-types in the mDIABLO model, but not in the standard DIABLO model (Fig. 4d). Tryptophan metabolism and Valine, leucine and isoleucine metabolism pathways were identified in both the gene and metabolite module datasets using mDIABLO. The heatmap of pairwise associations of all features identified with mDIABLO demonstrated the ability of DIABLO to select groups of correlated features which were predictive of pre- and post-challenge samples. The Asthma pathway was also identified [even though individual gene members were not significantly altered post-challenge (**Supplementary Fig. 10**)] and was negatively associated with Butanoate metabolism and positively associated with basophils, a hallmark cell-type in asthma (Fig. 4e). These findings depict DIABLO’s flexibility and sensitivity to detect subtle differences between repeated designs, and its ability to identify common molecular processes spanning different biological layers. The biological pathways identified suggest a mechanistic link with response to allergen challenge.

**Figure 4.**
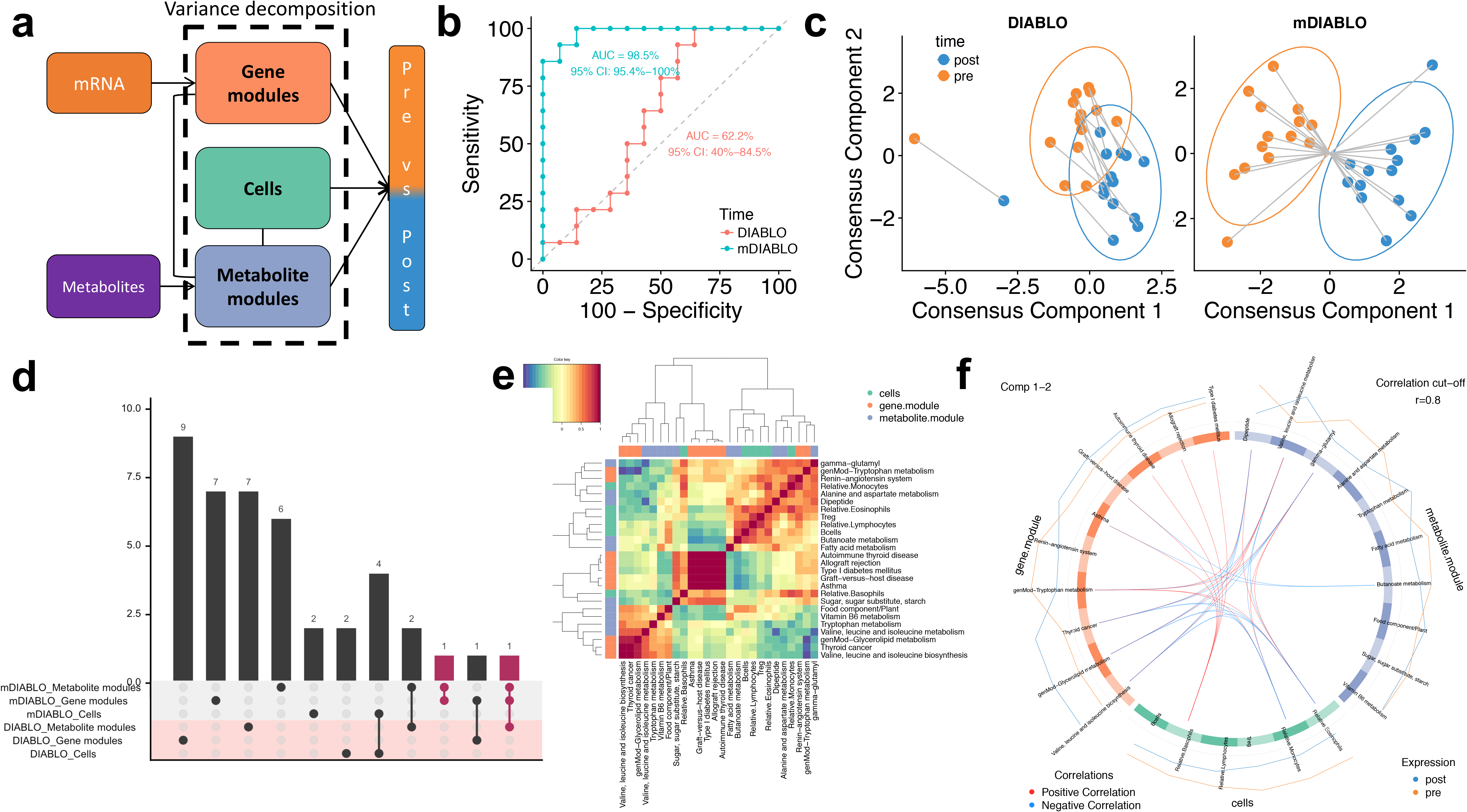
Asthma study: cross-over design and module-based analysis with DIABLO. **a)** DIABLO design includes a module-based decomposition approach to discriminate pre-and post-inhalation challenge samples. **b)** Receiver operating characteristic curves comparing the performance of the standard DIABLO and ‘multilevel DIABLO’ for repeated measures (mDIABLO) using leave-one-out cross-validation. **c)** Component plots depicting the separation of the pre- and post-challenge samples based on DIABLO and mDIABLO. **d)** Overlapping features selected from either DIABLO or mDIABLO. **e)** Heatmap of the Pearson correlation values between the features selected with mDIABLO. **f)** Circos plot depicting the strongest correlations between different omics features from the mDIABLO panel.

## Discussion

DIABLO aims to identify coherent patterns between datasets that change with respect different phenotypes. This purely data-driven, holistic, and hypothesis-free tool can be used to derive robust biomarkers and, ultimately, improve our understanding of the molecular mechanisms that drive disease.

We found that unsupervised methods identified features that formed strong interconnected multi-omics networks, but had poor discriminative ability. In contrast, features identified by supervised methods were discriminative, but formed sparsely connected networks. This trade-off between correlation and discrimination is a fundamental challenge when trying to identify biologically relevant biomarkers that are also clinically relevant [31]. DIABLO controls this trade-off by incorporating *a priori* relationships between different omic domains to adequately model dysregulated biological mechanisms between phenotypic conditions. This may explain the superior biological enrichment of the DIABLO_full models in our benchmarking experiments where the mRNA and miRNA expression as well as methylation activity were assumed to be correlated (Table 2). Since these omic domains are known to form real regulatory relationships in order to control complex biological processes, these multi-omic biomarker panels may be capturing this biological complexity. In contrast, these biomarkers were not uncovered when no association was assumed between omic datasets, as in the case of the DIABLO_null models and existing multi-step integrative strategies. Therefore, by controlling the trade-off between correlation and discrimination, DIABLO uncovered novel multi-omics biomarkers that have not previously been identified using existing integrative strategies. These novel biomarkers were part of densely connected clusters of omic variables which have prior known biological associations, further suggesting their potential biological plausibility.

There are areas of improvement that DIABLO will benefit from in the near future. The assumption of linear relationship between the selected omics features to explain the phenotypic response may not apply in some biological research areas, for example when integrating distance-based metagenomics studies, where kernel approaches could be further explored [32]. Selecting the optimal number of variables requires repeated cross-validation to ensure unbiased classification error rate evaluation. A grid approach was deemed reasonable and provided very good performance results, but several iterations to refine the grid may be required depending on the complexity of the classification problem. The grid search algorithm was recently improved [12], but we advise using a broad filtering strategy to alleviate computational time when dealing with extremely large datasets (e.g. > 50,000 features each). DIABLO was primarily developed for omics-measurements on a continuous scale after normalization, and further developments are needed for categorical data types, such as genotype data, as mentioned in [12]. Finally, DIABLO, like other methods we benchmarked, will be affected by technical artifacts of the data, such as batch effects and presence of confounding variables that may affect downstream integrative analyses. Therefore, we recommend exploratory analyses be carried out in each single omics dataset to assess the effect, if any, of technical factors and use of batch removal methods prior to the integration analysis [33–35].

To summarize, DIABLO is a versatile, component-based method that can integrate multiple high dimensional datasets and identify key variables that discriminate between phenotypic groups. DIABLO identified more biologically relevant and tightly correlated features across datasets when compared to existing multi-step classification schemes and integrative methods. The framework is highly flexible, suitable for single point or repeated measures study designs, and can accommodate various data transformations, such as feature summarization at the pathway level to enhance biological interpretability. DIABLO’s implementation includes intuitive graphical outputs to facilitate the interpretation of integrative analyses.

## Online Methods

### Code availability and software tool requirements

The DIABLO framework is implemented in the mixOmics R package [12]. mixOmics currently includes 19 multivariate methodologies, for single-omics and integrative analyses. All scripts and tutorials are provided in our companion web-page http://www.mixomics.org/mixDIABLO. All analyses were performed using the R statistical computing program (version 3.4.1) and the mixOmics package (version 6.3.0).

### Statistical methods and analysis

#### General multivariate framework to integrate multiple datasets measured on the same samples

DIABLO extends sparse generalized canonical correlation analysis (sGCCA) [15] to a classification (supervised) framework. sGCCA is a multivariate dimension reduction technique that uses singular value decomposition and selects co-expressed (correlated) variables from several omics datasets in a computationally and statistically efficient manner. sGCCA maximizes the covariance between linear combinations of variables (latent component scores) and projects the data into the smaller dimensional subspace spanned by the components. The selection of the correlated molecules across omics levels is performed internally in sGCCA with l_1_ –penalization on the variable coefficient vector defining the linear combinations. *Note that since all latent components are scaled in the algorithm, sGCCA maximizes the correlation between components. However, we will retain the term ‘covariance’ instead of ‘correlation’ throughout this section to present the general sGCCA framework*.

Denote *K* normalized, centered and scaled datasets *X_1_ (n* x *p_1_*), …, *X_K_* (*n* x *p_K_*), measuring the expression levels of *p_1,_ p_2, …,_ p_K_* omics variables on the same *n* samples, *k = 1, …, K*, sGCCA solves the optimization function:

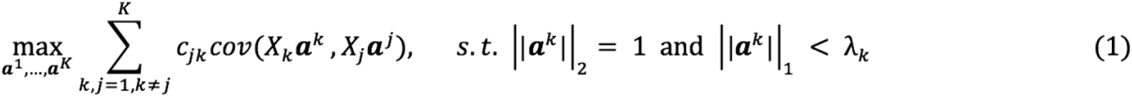

where *c_jk_* indicates whether to maximize the covariance between the datasets *X*_*k*_ and *X*_*j*_ according to the design matrix, with *c_jk_* values ranging from 0 (no relationship modelled between the datasets) to 1 otherwise, ***a***^*k*^ is the variable coefficient vector for each dataset *X*_*k*_, *λ*_k_ is a non-negative parameter that controls the amount of shrinkage and thus the number of non-zero coefficients in ***a***^*k*^. Similar to Lasso [36] and other l_1_ – penalized multivariate models developed for single omics analysis [14], the l_1_ penalization improves the interpretability of the component scores *X*_*k*_ ***a***^*k*^ that is now only defined on a subset of variables with non-zero coefficients in *X*_*k*_. The result is the identification of variables that are highly correlated between and within omics datasets.

Equation (1) describes the sGCCA model for the first dimension. Once the first set of coefficient vectors 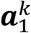 and associated component scores 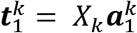 are obtained, residual matrices are calculated during the ‘deflation’ step for the second dimension, such that 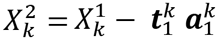, where 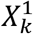 is the original centered and scaled data matrix. The subsequent set of components scores and coefficient vectors are then obtained by substituting *X*_*k*_ by 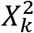 in (1). This process is repeated until a sufficient number of dimensions (or set of components) is obtained.

The underlying assumption of the sGCCA model is that the major source of common biological variation can be extracted via the first sets of component scores *X*_*k*_ ***a***^*k*^, while any unwanted variation due to heterogeneity across the datasets *X_K_* does not impact the statistical model. The optimization problem (1) is solved using a monotonically convergent algorithm [15].

#### DIABLO for supervised analysis and prediction

To extend sGCCA for a classification framework, we substitute one omics dataset *X_k_* in (1) with a dummy indicator matrix *Y* of size (*n* x *G*), where *G* is the number of phenotype groups that indicate the class membership of each sample. In addition, and for easier use of the method, we replaced the l_1_ penalty parameter *λ*_k_ by the number of variables to select in each dataset and each component, as there is a direct correspondence between both parameters.

Denote a new sample *i* which is measured across the different types of omics datasets 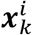, its class membership is predicted by the fitted sGCCA model with the estimated variable coefficients vectors 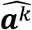 to obtain the predicted scores 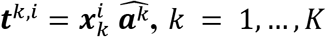. Therefore, to each dataset *k* corresponds a predicted continuous score ***t***^*k*,*i*^. The predicted class of sample *i* for each dataset is obtained from the predicted score using one of the distances Maximum, Centroids or Mahalanobis [37] as described in [12]. The consensus class membership is determined using either a majority vote, or by averaging all ***t***^*k*,*i*^ across all *K* datasets before using the prediction distance of choice (‘average prediction’ scheme). In case of ties in the majority vote scheme, ‘NA’ is allocated as a prediction but is counted as a misclassification error during the performance evaluation. As the class prediction relies on individual vote from each omics set, DIABLO allows for some missing datasets *X*_*k*_ during the prediction step, as illustrated in the Breast Cancer case study. We used the centroid distance for the weighted majority vote scheme (breast cancer study) and the maximum distance for the average vote scheme (asthma study) as those led to best performance (see [12] for details about distance measures and voting schemes that can be used).

#### Design matrix in DIABLO

The design matrix *C* is a (*K* x *K*) matrix with values ranging from 0 to 1 which specifies whether the covariance between two datasets should be maximized DIABLO (see equation (1)). In our simulation study, we evaluated two scenarios: a null design (DIABLO_null) when no omics datasets are connected, and a full design when all datasets are connected (DIABLO_full):

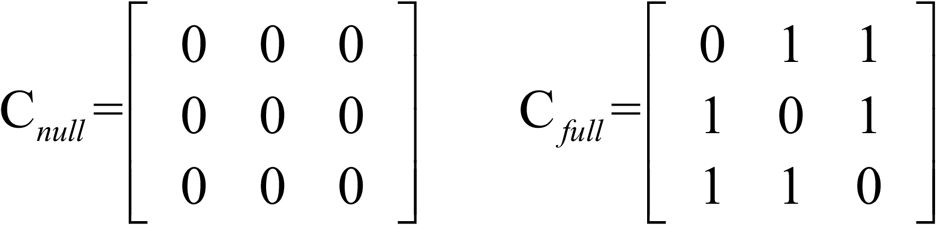

However, every dataset is connected to the outcome *Y* internally in the method. For the two case studies (breast cancer and asthma) the design matrix was chosen based on our proposed method (see below ***Parameters tuning***). Note that the design matrix is not restricted to 0 and 1 values only and a compromise between correlation and discrimination can also be modelled as described in [12].

#### Input data in DIABLO

While DIABLO does not assume particular data distributions, all datasets should be normalized appropriately according to each omics platform and preprocessed if necessary (see normalization steps described below for each case study). Samples should be represented in rows in the data matrices and match the same sample across omics datasets. The phenotype outcome *Y* is a factor indicating the class membership of each sample. The R function in mixOmics will internally center and scale each variable as is conventionally performed in PLS-based models and will create the dummy matrix outcome from Y. A multilevel variance decomposition option is available for repeated measures study designs (see below).

#### Parameters tuning

The first parameter to tune is the design matrix C, which can be determined using either prior biological knowledge, or a data-driven approach. The latter approach uses PLS method implemented in mixOmics that models pair-wise associations between omics datasets. If the correlation between the first component of each omics dataset is above a given threshold (e.g. 0.8) then a connection between those datasets is included in the DIABLO design as a 1 value.

The second parameter to tune is the total number of components. In several analyses we found that G − 1 components were sufficient to extract sufficient information to discriminate all phenotype groups [14], but this can be assessed by evaluating the model performance across all specified components (described below) as well as using graphical outputs such as sample plots to visualize the discriminatory ability of each component.

Finally, the third set of parameters to tune is the number of variables to select per dataset and per component. Such tuning can rapidly become cumbersome, as there might be numerous combinations of selection sizes to evaluate across all *K* datasets. For the breast cancer study, we used 5-fold cross-validation repeated 50 times to evaluate the performance of the model over a grid of different possible values of variables to select (**Supplementary Fig. 8**). The performance of the model for a given set of parameters (including number of component and number of variables to select) was based on the balanced classification error rate using majority vote or average prediction schemes with centroids distance. The balanced classification error rate is useful in the case of imbalanced class sizes, where the majority classes can have strong influence on the overall error rate. The balanced error rate measure calculates the weighted average of the individual class error rates with respect to their class sample size. In our experience, the number of variables to select in each dataset provided less of an improvement on the error rate compared to tuning the number of components. Therefore, even a grid composed of a small number of variables (<50 with steps of 5 or 10) may suffice as it does not substantially change the classification performance. This is because of the use of regularization constraints which reduces the variability in the variable coefficients and thus maintains the predictive ability of the model. Further, the variable selection size can also be guided according to the downstream biological interpretation to be performed. For example, a gene-set enrichment analysis may require a larger set of features than a literature-search interpretation.

#### Visualization outputs with DIABLO

To facilitate the interpretation of the integrative analysis, several types of graphical outputs were implemented in mixOmics.

#### Sample plots

The consensus plot which depicts the samples is computed by calculating the average of the components from each dataset. Omics specific samples plots can also be obtained by plotting components associated to each data set. The sample plot are useful to visualize the ability of the DIABLO model to extract common information at the sample level for each dataset, and the discriminatory power of each data type to separate the phenotypic groups. The scatterplot matrix represents the correlation between components for the same dimension but across all omics datasets. This plot assesses the model’s ability to maximize the correlation as indicated in the design matrix. Separation of subjects according to their phenotypic groups can be visualized.

#### Variable plots

To visualize selected variables, we proposed circos plot to represent correlations between and within variables from each dataset at the variable level. The association between variables is computed using a similarity score that is analogous to a Pearson correlation coefficient, as previously described in [38]. For each omics dataset, DIABLO produces a variable coefficient matrix of size (*p*_*k*_X*H*), where *H* is the total number of components in the model. The product of any two matrices approximates the association score between variables of the two omics datasets. The association between variables is displayed as a color-coded link inside the plot to represent a positive or negative correlation above a user-specified threshold. The selected variables are represented on the side of the circos plot, with side colors indicating each omics type, optional line plots represent the expression levels in each phenotypic group.

#### Clustered Image Map (CIM)

A clustered image map [38] based on the Euclidean distance and the complete linkage displays an unsupervised clustering between the selected variables (centered and scaled) and the samples. Color bars represent the sample phenotypic groups (columns) and the type of omics (rows) variables.

### Gene-set enrichment analyses

Significance of enrichment was determined using a hypergeometric test of the overlap between the selected features (mapped to official HUGO gene symbols or official miRNA symbols) and the various gene sets contained in the collections. In order to carry out the comparison, each feature set was mapped back to official HUGO gene symbols. This was done as follows across the respective data types: mRNA, CpGs and proteins. The following collections were used as gene-sets for the enrichment analysis [39]: C1 - positional gene sets for each human chromosome and cytogenetic band. C2 – curated gene sets (Pathway Interaction DB [PID], Biocarta [BIOCARTA], Kyoto Encyclopedia of Genes and Genomes [KEGG], Reactome [REACTOME], and others), C3 - motif gene sets based on conserved cis-regulatory motifs from a comparative analysis of the human, mouse, rat, and dog genomes. C4 – computational gene sets (from the Cancer Gene Neighbourhoods [CGN] and Cancer Modules [CM] – citation available via the MolSigDB [23]. C5 - GO gene sets consist of genes annotated by the same GO terms. C6 –ontologic gene sets (Gene sets represent signatures of cellular pathways which are often dis-regulated in cancer). C7 - immunologic gene sets defined directly from microarray gene expression data from immunologic studies. H - hallmark gene sets are coherently expressed signatures derived by aggregating many MSigDB gene sets to represent well-defined biological states or processes. & A. BTM - Blood Transcriptional Modules [40]. B. TISSUES - cell-specific expression from Benita *et al.* [41].

#### Modular analysis

Eigengene summarization is a common approach to decompose a *n* x *p* dataset (where *n* is the number of samples and *p* is the number of variables in a module), to a component (linear combination of all *p* variables) that represents the summarized expression of genes in the module [18]. For the asthma study, 15,683 genes were reduced to 229 KEGG pathways and 292 metabolites were reduced to 60 metabolic pathways using eigengene summarization.

#### Multilevel transformation

For multivariate analyses, A multilevel approach separates the within subject variation matrix (*X_w_*) and the between subject variation (*X_b_*) for a given dataset (*X*) [42], ie. *X = X_w_ + X_b_*. In the case of a two-repeated measured problem (e.g. pre vs post challenge), the within subject variation matrix is similar to calculating the net difference for each individual between the data obtained for pre and post challenge. For each omics dataset, the within-subject variation matrix was extracted prior to applying DIABLO. In the asthma study, the multilevel approach (called variance decomposition step) was applied to the cell-type, gene and metabolite module datasets.

## Declarations

### Acknowledgements

The authors would like to thank Mr Kevin Chang (University of Auckland) for some preliminary exploratory analyses of the breast cancer dataset. We would also like to thank Dr Chao Liu (University of Queensland) for obtaining the PAM50 phenotypic information for the TCGA datasets.

### Authors’ contributions

AS performed the data pre-processing, the statistical analyses and developed the DIABLO method. BG implemented the R scripts for DIABLO and graphical outputs, CPS performed the gene enrichment analyses, MV implemented the circos plots, FR and BG implemented the R scripts in mixOmics along with the S3 functions, SJT supervised AS and participated in the design of the study. KALC supervised AS, BG, MV and FR, participated in the development of the DIABLO method and provided statistical advice. AS and KALC edited the manuscript, with editorial input from SJT and CPS.

## References

1. Zhu J, Sova P, Xu Q, Dombek KM, Xu EY, Vu H, et al. Stitching together multiple data dimensions reveals interacting metabolomic and transcriptomic networks that modulate cell regulation. Levchenko A, editor. PLoS Biol [Internet]. 2012 [cited 2016 Jan 19];10:e1001301. Available from: http://dx.plos.org/10.1371/journal.pbio.1001301

2. Kim D, Li R, Dudek SM, Ritchie MD. ATHENA: Identifying interactions between different levels of genomic data associated with cancer clinical outcomes using grammatical evolution neural network. BioData Min. 2013;6:23.

3. Wang B, Mezlini AM, Demir F, Fiume M, Tu Z, Brudno M, et al. Similarity network fusion for aggregating data types on a genomic scale. Nat Methods [Internet]. 2014 [cited 2016 Jan 19];11:333–7. Available from: http://www.nature.com/doifinder/10.1038/nmeth.2810

4. Ritchie MD, Holzinger ER, Li R, Pendergrass SA, Kim D. Methods of integrating data to uncover genotype–phenotype interactions. Nat Rev Genet [Internet]. 2015 [cited 2015 Jul 10];16:85–97. Available from: http://www.nature.com/doifinder/10.1038/nrg3868

5. Yugi K, Kubota H, Hatano A, Kuroda S. Trans-Omics: How To Reconstruct Biochemical Networks Across Multiple ‘Omic’ Layers. Trends Biotechnol [Internet]. 2016 [cited 2018 Feb 21];34:276–90. Available from: http://linkinghub.elsevier.com/retrieve/pii/S0167779915002735

6. Günther O, Chen V, Freue GC, Balshaw R, Tebbutt S, Hollander Z, et al. A computational pipeline for the development of multi-marker bio-signature panels and ensemble classifiers. 2012 [cited 2016 Jan 19];13:326. Available from: http://summit.sfu.ca/item/13303

7. Aben N, Vis DJ, Michaut M, Wessels LFA. TANDEM: a two-stage approach to maximize interpretability of drug response models based on multiple molecular data types. Bioinformatics [Internet]. 2016 [cited 2017 Aug 2];32:i413–20. Available from: https://academic.oup.com/bioinformatics/article-lookup/doi/10.1093/bioinformatics/btw449

8. Ma S, Ren J, Fenyö D. Breast cancer prognostics using multi-omics data. AMIA Summits Transl Sci Proc [Internet]. 2016 [cited 2017 May 30];2016:52. Available from: https://www.ncbi.nlm.nih.gov/pmc/articles/PMC5001766/

9. Bersanelli M, Mosca E, Remondini D, Giampieri E, Sala C, Castellani G, et al. Methods for the integration of multi-omics data: mathematical aspects. BMC Bioinformatics [Internet]. 2016 [cited 2016 May 8];17. Available from: http://www.biomedcentral.com/1471-2105/17/S2/15

10. Meng C, Zeleznik OA, Thallinger GG, Kuster B, Gholami AM, Culhane AC. Dimension reduction techniques for the integrative analysis of multi-omics data. Brief Bioinform [Internet]. 2016 [cited 2018 Feb 21];17:628–41. Available from: https://academic.oup.com/bib/article-lookup/doi/10.1093/bib/bbv108

11. Huang S, Chaudhary K, Garmire LX. More Is Better: Recent Progress in Multi-Omics Data Integration Methods. Front Genet [Internet]. 2017 [cited 2018 Feb 21];8. Available from: http://journal.frontiersin.org/article/10.3389/fgene.2017.00084/full

12. Rohart F, Gautier B, Singh A, Cao K-AL. mixOmics: An R package for ‘omics feature selection and multiple data integration. PLOS Comput Biol [Internet]. 2017 [cited 2018 Jan 29];13:e1005752. Available from: http://journals.plos.org/ploscompbiol/article?id=10.1371/journal.pcbi.1005752

13. Wold H. Estimation of Principal Components and Related Models by Iterative Least squares. Multivar Anal. 1966;391–420.

14. Lê Cao K-A, Boitard S, Besse P. Sparse PLS discriminant analysis: biologically relevant feature selection and graphical displays for multiclass problems. BMC Bioinformatics [Internet]. 2011 [cited 2015 Jul 15];12:253. Available from: http://www.biomedcentral.com/1471-2105/12/253/

15. Tenenhaus A, Philippe C, Guillemot V, Le Cao K-A, Grill J, Frouin V. Variable selection for generalized canonical correlation analysis. Biostatistics [Internet]. 2014 [cited 2015 Jul 15];15:569–83. Available from: http://biostatistics.oxfordjournals.org/cgi/doi/10.1093/biostatistics/kxu001

16. Witten DM, Tibshirani R, Hastie T. A penalized matrix decomposition, with applications to sparse principal components and canonical correlation analysis. Biostatistics [Internet]. 2009 [cited 2016 Jul 27];10:515–34. Available from: http://biostatistics.oxfordjournals.org/cgi/doi/10.1093/biostatistics/kxp008

17. Lee HK, Hsu AK, Sajdak J, Qin J, Pavlidis P. Coexpression analysis of human genes across many microarray data sets. Genome Res [Internet]. 2004 [cited 2016 Mar 30];14:1085–1094. Available from: http://genome.cshlp.org/content/14/6/1085.short

18. Langfelder P, Horvath S. WGCNA: an R package for weighted correlation network analysis. BMC Bioinformatics [Internet]. 2008 [cited 2016 Apr 4];9:559. Available from: http://www.biomedcentral.com/1471-2105/9/559

19. The TCGA Research Network. The Cancer Genome Atlas [Internet]. Available from: http://cancergenome.nih.gov/

20. Singh A, Yamamoto M, Kam SHY, Ruan J, Gauvreau GM, O’Byrne PM, et al. Gene-metabolite expression in blood can discriminate allergen-induced isolated early from dual asthmatic responses. Hsu Y-H, editor. PLoS ONE [Internet]. 2013 [cited 2015 Jul 18];8:e67907. Available from: http://dx.plos.org/10.1371/journal.pone.0067907

21. Singh A, Yamamoto M, Ruan J, Choi JY, Gauvreau GM, Olek S, et al. Th17/Treg ratio derived using DNA methylation analysis is associated with the late phase asthmatic response. Allergy Asthma Clin Immunol [Internet]. 2014 [cited 2016 Mar 2];10:32. Available from: http://www.biomedcentral.com/content/pdf/1710-1492-10-32.pdf

22. Lock EF, Hoadley KA, Marron JS, Nobel AB. Joint and individual variation explained (JIVE) for integrated analysis of multiple data types. Ann Appl Stat [Internet]. 2013 [cited 2018 Jan 24];7:523–42. Available from: http://projecteuclid.org/euclid.aoas/1365527209

23. Liberzon A, Birger C, Thorvaldsdóttir H, Ghandi M, Mesirov JP, Tamayo P. The Molecular Signatures Database Hallmark Gene Set Collection. Cell Syst [Internet]. 2015 [cited 2018 Jan 30];1:417–25. Available from: http://linkinghub.elsevier.com/retrieve/pii/S2405471215002185

24. Xie B, Ding Q, Han H, Wu D. miRCancer: a microRNA-cancer association database constructed by text mining on literature. Bioinformatics [Internet]. 2013 [cited 2018 Jan 30];29:638–44. Available from: https://academic.oup.com/bioinformatics/article-lookup/doi/10.1093/bioinformatics/btt014

25. Hamosh A. Online Mendelian Inheritance in Man (OMIM), a knowledgebase of human genes and genetic disorders. Nucleic Acids Res [Internet]. 2004 [cited 2018 Jan 30];33:D514–7. Available from: https://academic.oup.com/nar/article-lookup/doi/10.1093/nar/gki033

26. Chung I-F, Chen C-Y, Su S-C, Li C-Y, Wu K-J, Wang H-W, et al. DriverDBv2: a database for human cancer driver gene research. Nucleic Acids Res [Internet]. 2016 [cited 2018 Jan 30];44:D975–9. Available from: https://academic.oup.com/nar/article-lookup/doi/10.1093/nar/gkv1314

27. Liquet B, Lê Cao K-A, Hocini H, Thiébaut R. A novel approach for biomarker selection and the integration of repeated measures experiments from two assays. BMC Bioinformatics [Internet]. 2012 [cited 2015 Jul 18];13:325. Available from: http://www.biomedcentral.com/1471-2105/13/325/

28. Allahyar A, de Ridder J. FERAL: network-based classifier with application to breast cancer outcome prediction. Bioinformatics [Internet]. 2015 [cited 2018 Feb 1];31:i311–9. Available from: https://academic.oup.com/bioinformatics/article-lookup/doi/10.1093/bioinformatics/btv255

29. Cun Y, Fröhlich H. Network and data integration for biomarker signature discovery via network smoothed t-statistics. Boccaletti S, editor. PLoS ONE [Internet]. 2013 [cited 2017 May 30];8:e73074. Available from: http://dx.plos.org/10.1371/journal.pone.0073074

30. Sokolov A, Carlin DE, Paull EO, Baertsch R, Stuart JM. Pathway-based genomics prediction using generalized elastic net. PLoS Comput Biol [Internet]. 2016 [cited 2017 May 30];12:e1004790. Available from: http://journals.plos.org/ploscompbiol/article?id=10.1371/journal.pcbi.1004790

31. Wang TJ. Assessing the Role of Circulating, Genetic, and Imaging Biomarkers in Cardiovascular Risk Prediction. Circulation [Internet]. 2011 [cited 2018 Feb 23];123:551–65. Available from: http://circ.ahajournals.org/cgi/doi/10.1161/CIRCULATIONAHA.109.912568

32. Mariette J, Villa-Vialaneix N. Unsupervised multiple kernel learning for heterogeneous data integration. Bioinformatics [Internet]. 2017 [cited 2018 Mar 6]; Available from: http://academic.oup.com/bioinformatics/advance-article/doi/10.1093/bioinformatics/btx682/4565592

33. Johnson WE, Li C, Rabinovic A. Adjusting batch effects in microarray expression data using empirical Bayes methods. Biostatistics [Internet]. 2007 [cited 2016 May 12];8:118–27. Available from: http://biostatistics.oxfordjournals.org/cgi/doi/10.1093/biostatistics/kxj037

34. Gagnon-Bartsch JA, Speed TP. Using control genes to correct for unwanted variation in microarray data. Biostatistics [Internet]. 2012 [cited 2018 Mar 6];13:539–52. Available from: https://academic.oup.com/biostatistics/article-lookup/doi/10.1093/biostatistics/kxr034

35. Parker HS, Corrada Bravo H, Leek JT. Removing batch effects for prediction problems with frozen surrogate variable analysis. PeerJ [Internet]. 2014 [cited 2016 May 12];2:e561. Available from: https://peerj.com/articles/561

36. Tibshirani R. Regression shrinkage and selection via the lasso. J R Stat Soc Ser B Methodol. 1996;58:267–88.

37. Le Cao K-A, Gonzalez I, Dejean S. integrOmics: an R package to unravel relationships between two omics datasets. Bioinformatics [Internet]. 2009 [cited 2016 Apr 3];25:2855–6. Available from: http://bioinformatics.oxfordjournals.org/cgi/doi/10.1093/bioinformatics/btp515

38. González I, Lê Cao K-A, Davis MJ, Déjean S. Visualising associations between paired ‘omics’ data sets. BioData Min [Internet]. 2012 [cited 2015 Jul 15];5:1–23. Available from: http://link.springer.com/article/10.1186/1756-0381-5-19

39. Subramanian A, Tamayo P, Mootha VK, Mukherjee S, Ebert BL, Gillette MA, et al. Gene set enrichment analysis: a knowledge-based approach for interpreting genome-wide expression profiles. Proc Natl Acad Sci [Internet]. 2005 [cited 2016 Jul 26];102:15545–15550. Available from: http://www.pnas.org/content/102/43/15545.short

40. Chaussabel D, Quinn C, Shen J, Patel P, Glaser C, Baldwin N, et al. A Modular Analysis Framework for Blood Genomics Studies: Application to Systemic Lupus Erythematosus. Immunity [Internet]. 2008 [cited 2016 Jul 22];29:150–64. Available from: http://linkinghub.elsevier.com/retrieve/pii/S1074761308002835

41. Benita Y, Cao Z, Giallourakis C, Li C, Gardet A, Xavier RJ. Gene enrichment profiles reveal T-cell development, differentiation, and lineage-specific transcription factors including ZBTB25 as a novel NF-AT repressor. Blood [Internet]. 2010 [cited 2018 Mar 5];115:5376–84. Available from: http://www.bloodjournal.org/cgi/doi/10.1182/blood-2010-01-263855

42. Westerhuis JA, van Velzen EJJ, Hoefsloot HCJ, Smilde AK. Multivariate paired data analysis: multilevel PLSDA versus OPLSDA. Metabolomics [Internet]. 2010 [cited 2016 Jul 27];6:119–28. Available from: http://link.springer.com/10.1007/s11306-009-0185-z

