## Supplementary Materials for "DIABLO: from multi-omics assays to biomarker discovery, an integrative approach"

#### Supplementary Figures

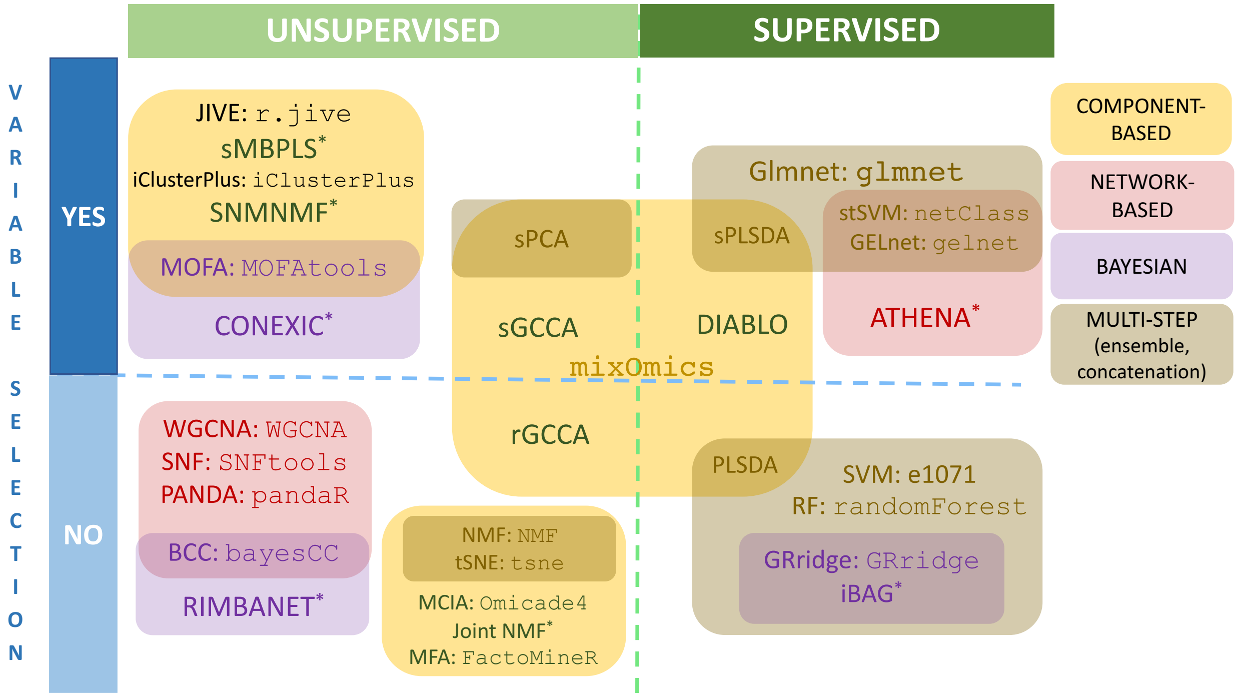

##### Supplementary Figure 1

Overview of approaches used for the integration of multiple high dimensional omics datasets using either unsupervised or supervised analyses. Most integrative methods were developed for unsupervised analyses. Variable selection is an important feature of the methods to improve interpretation of these complex models. Various types of integrative methods are listed, ranging from Component-based that reduce the dimensionality of high-throughput omics datasets, Bayesian methods, Network-based and multi-step approaches which include concatenation and ensemble approaches[1]. Concatenation-based approach combine multiple matrices and apply standard single omics analysis without taking into account the type of omics variable in the model. Ensemble-based approaches involve the development of independent models for each omics dataset, after which the outputs are combined using various voting schemes (e.g. majority vote, average vote). Methods name in courier font indicate the name of the R package. *Methods are coded in other languages are indicated below.

Abbreviations: JIVE: Joint and Individual Variation Explained[6], *sMBPLS: sparse Multiblock Partial Least Squares (Matlab)[7], SNMNMF: Sparse Network-regularized Multiple Non-negative Matrix Factorization (Matlab)[8], MOFA: Multi-Omics Factor Analysis[9], *CONEXIC: Copy Number and Expression In Cancer (Java)[10], WGCNA: Weighted Gene Co-expression Network Analysis[11], SNF: Similarity Network Fusion[12], PANDA: Passing Attributes between Networks for Data Assimilation[13], BCC: Bayesian Consensus Clustering[14], *RIMBANET: Reconstructing Integrative Molecular Bayesian Networks (Perl)[3]; sPCA : sparse Principal Component Analysis[15]; sGCCA: sparse generalized canonical correlation analysis[16]; rGCCA : regularized generalized canonical correlation analysis[17]; NMF: Non-Negative Factorization (Matlab); MFA: Multiple Co-inertia Analysis (MCIA); Multiple Factor Analysis[19]; glmnet: Lasso and Elastic-Net Regularized Generalized Linear Models[20]; sPLSDA: sparse Partial Least Squares Discriminant Analysis[21]; stSVM Smoothed t-statistics Support Vector Machine[22]; GELnet: Generalized Elastic Net[23]; *ATHENA: Analysis Tool for Heritable and Environmental Network Associations (Perl)[4]; SVM: Support Vector Machine; RF: Random Forest[24]; GRridge: Adaptive group-regularized ridge regression[25]; *iBAG: integrative Bayesian Analysis of Genomics (R and Shiny)[5]

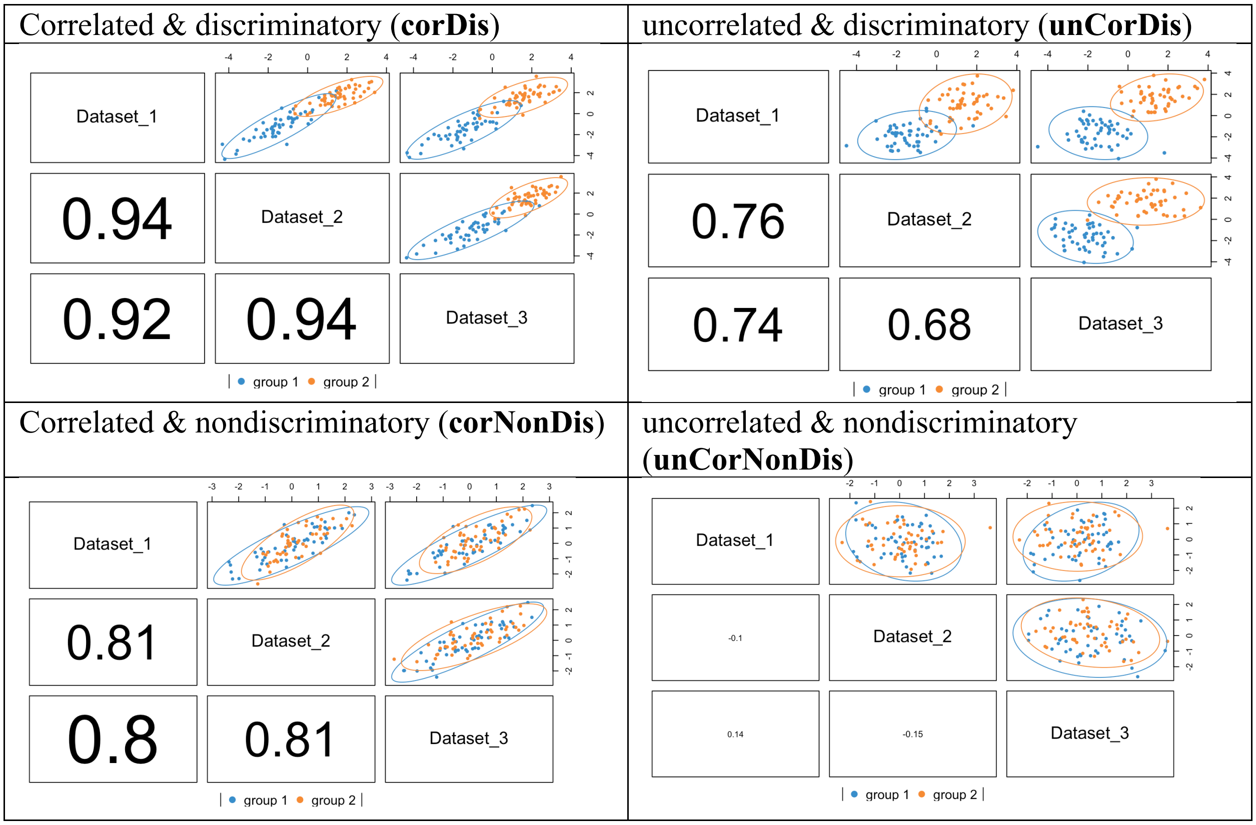

Supplementary Figure 2

**Simulation study.** Scatterplot matrices depicting the Pearon's correlation between the first components generated different types of omics variables. Different simulation scenarios where considered. Top left: strong correlation across multi-omics datasets and strong discrimination between phenotypic groups, as indicated by the high correlation coefficients. Top right: weak correlation across multi-omics datasets but strong discrimination between phenotypic groups as indicated by the clusters of samples belonging to either group 1 or group 2. Bottom left: strong correlation across multi- omics datasets but poor discrimination between phenotypic groups. Bottom right: weak correlation across multi-omics datasets and poor discrimination between phenotypic groups.

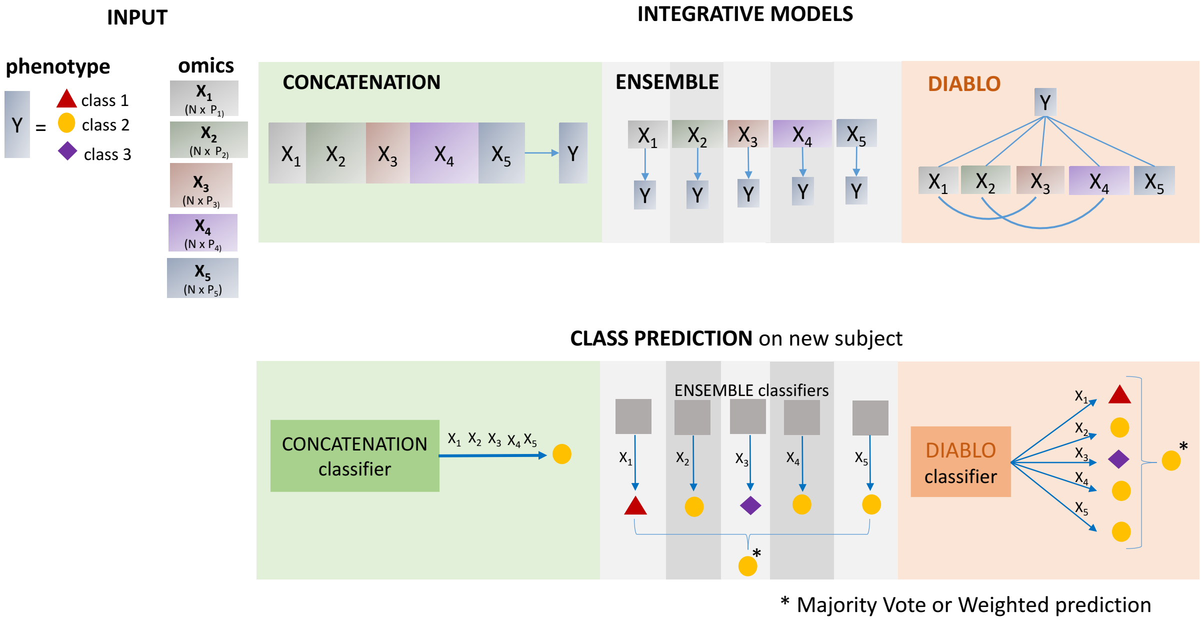

Supplementary Figure 3

**Integrative prediction frameworks including multi-step approaches (concatenation, ensemble) and DIABLO to identify multi-omics molecular signatures**. Concatenation-based integration combines multiple datasets into a single large dataset, with the aim to predict a phenotype of interest. Ensemble-based classification methods construct a predictive model on each individual dataset before combining the model predictions. None of these approaches account or model relationships between datasets and thus limit our understanding of molecular interactions at multiple functional levels. DIABLO simultaneously maximizes the associations between datasets and a phenotype of interest to identify a correlated set of variables of different omics-types that are also discriminatory. The prediction is based on each omics-associated component derived from the model (see **Supplementary Note**). All methods presented here are data-driven approaches, which do not use any prior knowledge such as from curated biological databases (eg. protein-protein interactions).

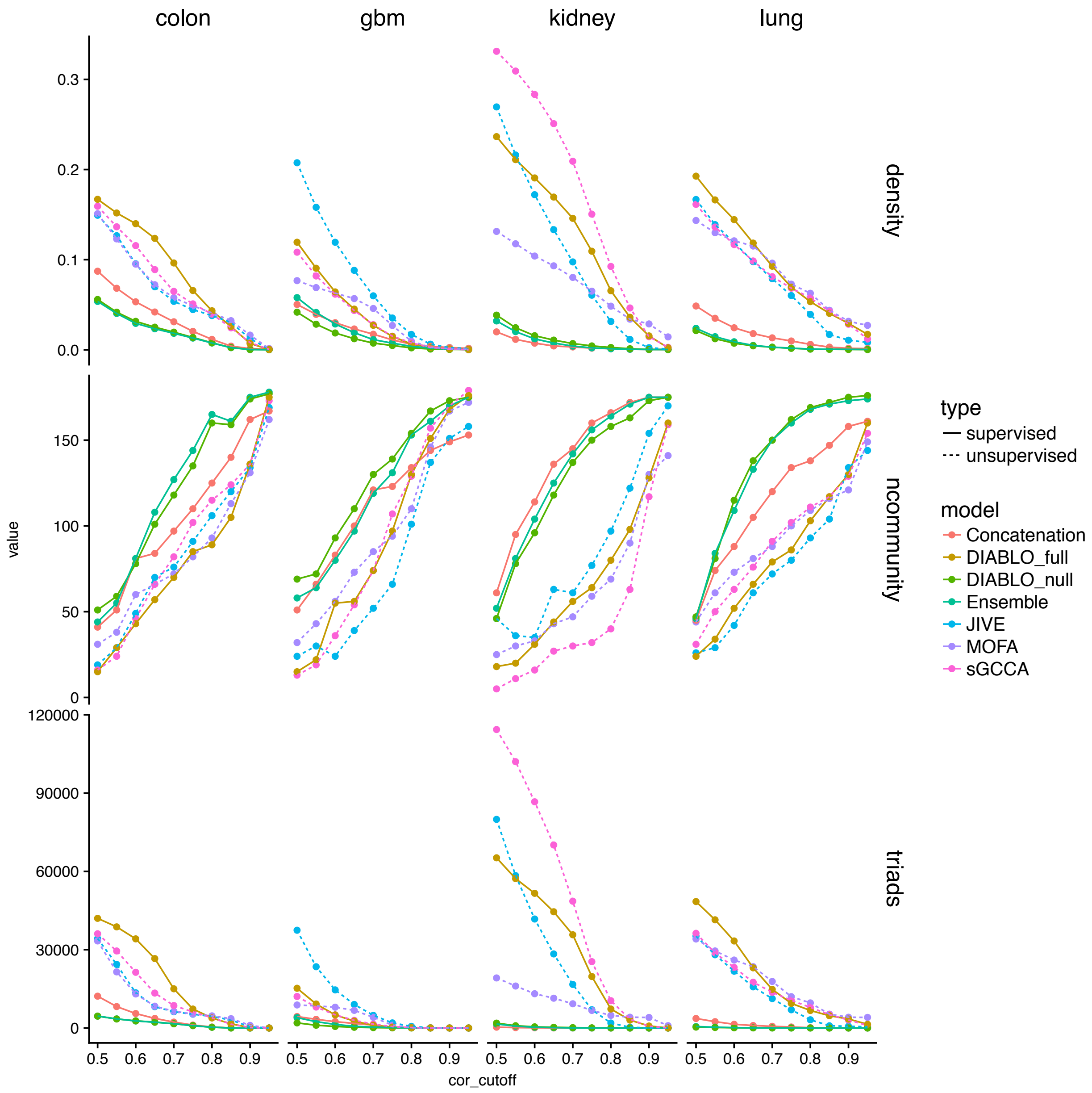

Supplementary Figure 4

**Benchmark analyses: network properties of multi-omics signatures.** We analysed each of the four multi-omics cancer datasets with component-based integrative methods with variable selection. The network attributes, density, number of communities and triads resulting from each molecular signature are represented. The unsupervised methods (dashed lines) led multi-omics signatures with a higher graph density, a greater number of triads and a lower number of communities as compared to supervised methods (solid lines), with the exception of DIABLO_full which simultaneously explains the correlation structure between multiple omic datasets and a phenotypic response variable.

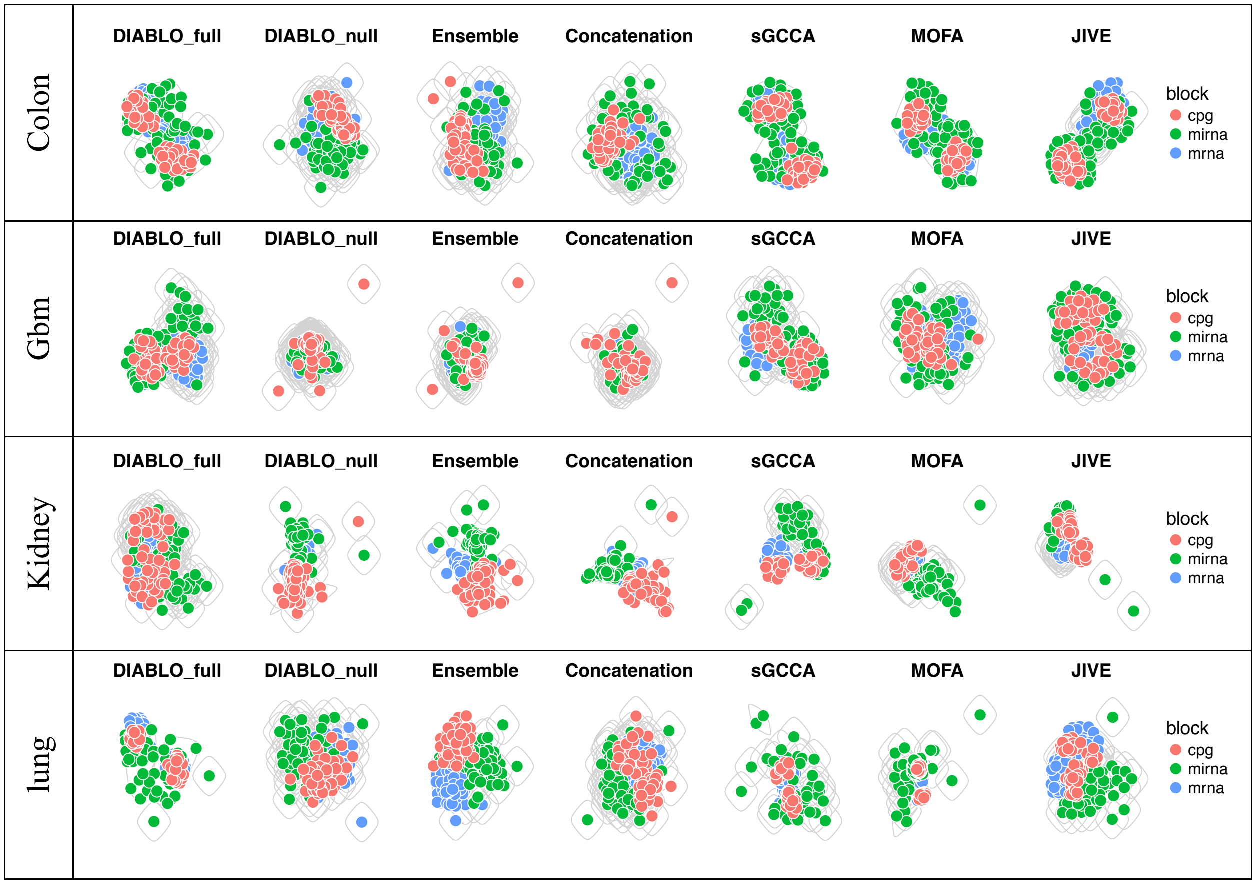

Supplementary Figure 5

**Benchmark analyses: network connectivity of multi-omics signatures.** Networks of the multi-omics biomarker panels identified from each method are represented for a Pearson's correlation cut-off of |0.4|.

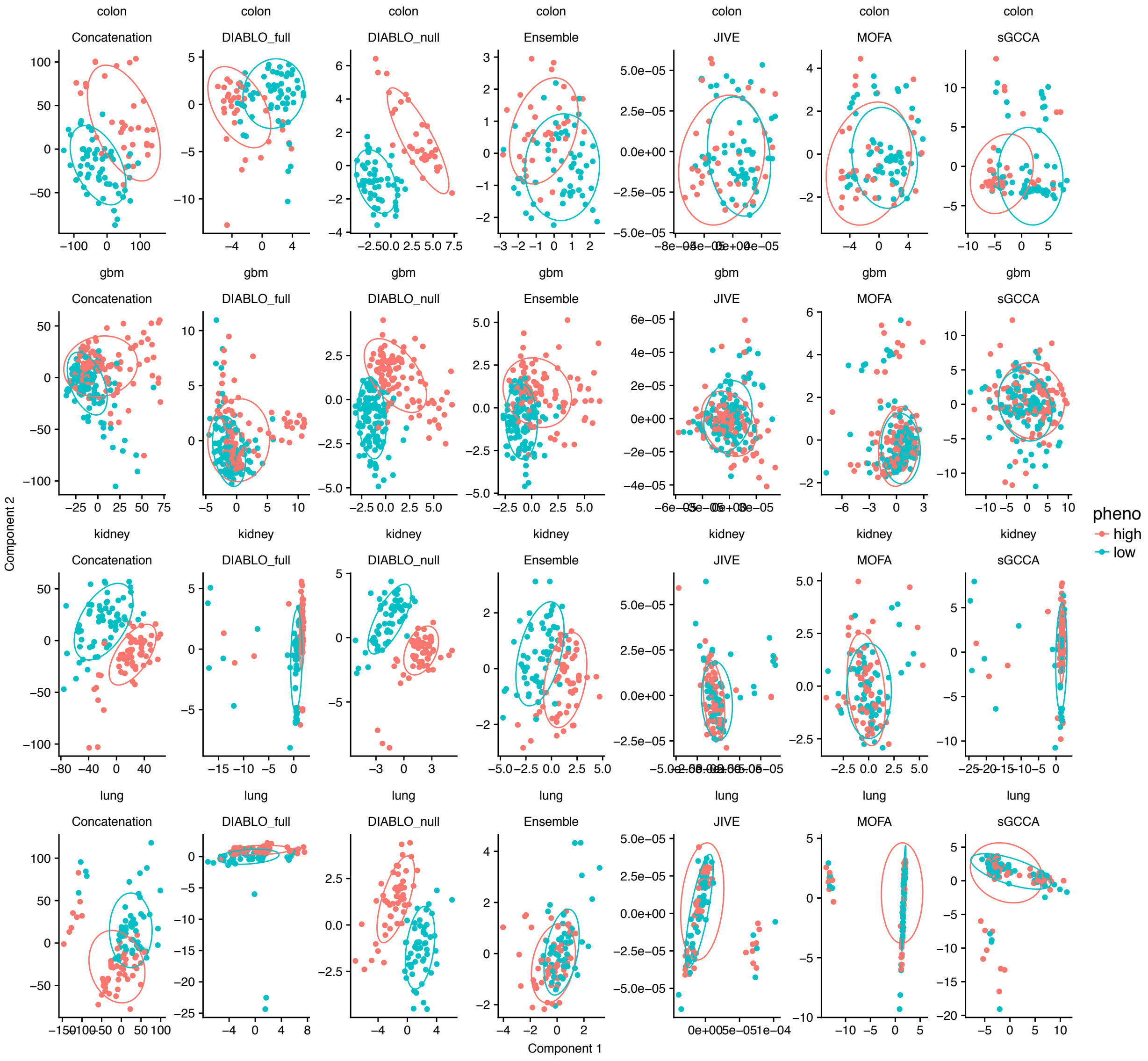

Supplementary Figure 6

**Benchmark analyses: sample plots for each multi-omics panel**. As expected, a strong separation between high and low survival groups can be observed for supervised methods but not for unsupervised methods. The level of discrimination decreases when using DIABLO_full compared to DIABLO_null.

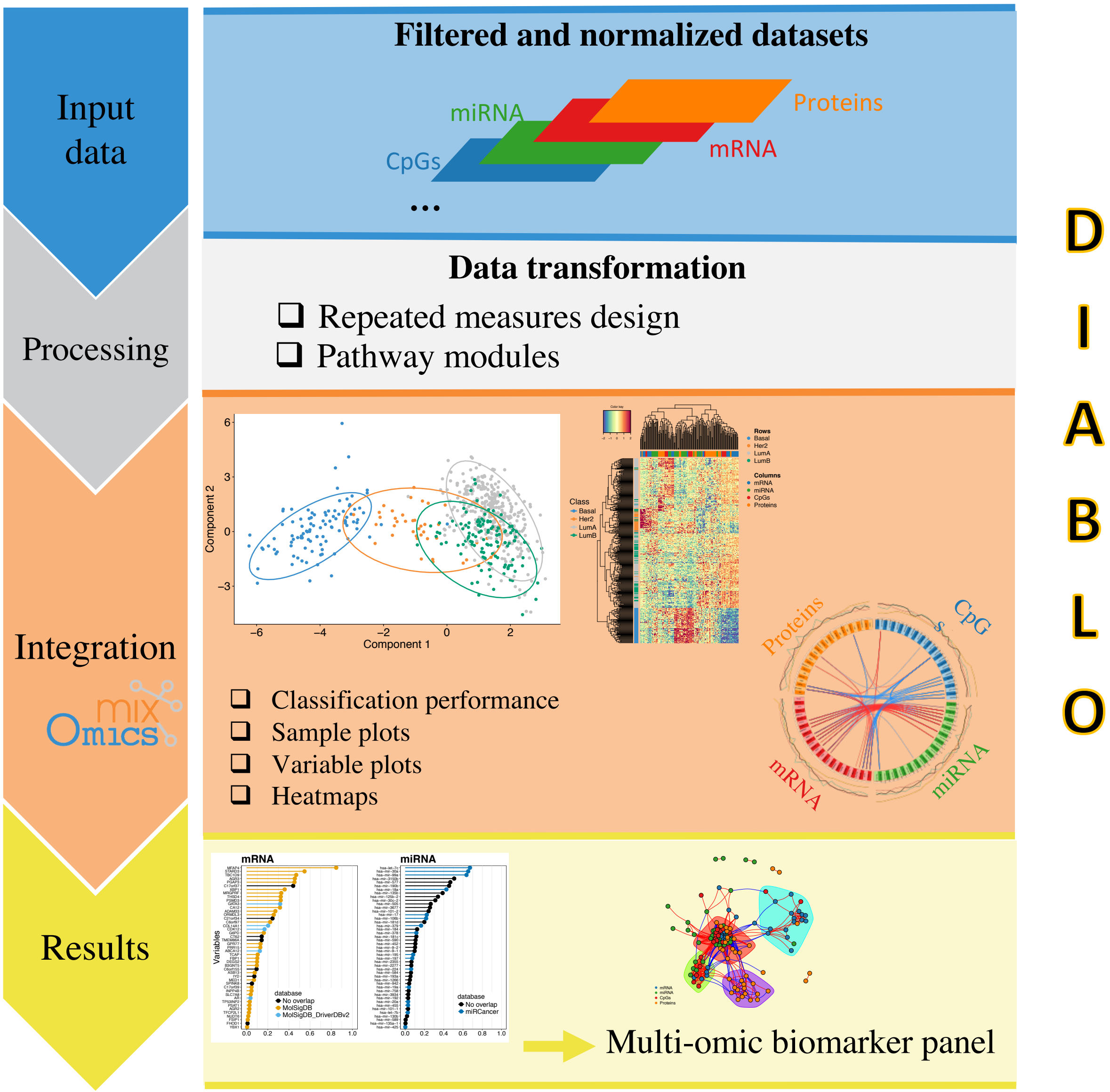

Supplementary Figure 7

**A standard DIABLO workflow.** The first step inputs multiple omics datasets measured on the same individuals, that were previously normalized and filtered, , along with the phenotype information indicating the class membership of each sample (two or more groups). Optional preprocessing steps include multilevel transformation for repeated measures study designs and pathway module summary transformations. DIABLO is a multivariate dimension reduction method that seeks for latent components – linear combinations of variables from each omics dataset, that are maximally correlated as specified by a design matrix (see Methods section). The identification of a multi-omics panel is obtained with $l_1$ penalties in the model that shrink the variable coefficients defining the components to zero. Numerous visualizations are proposed to provide insights into the multi-omics panel and guide the interpretation of the selected omics variables, including sample and variable plots. Downstream analysis include gene set enrichment analysis.

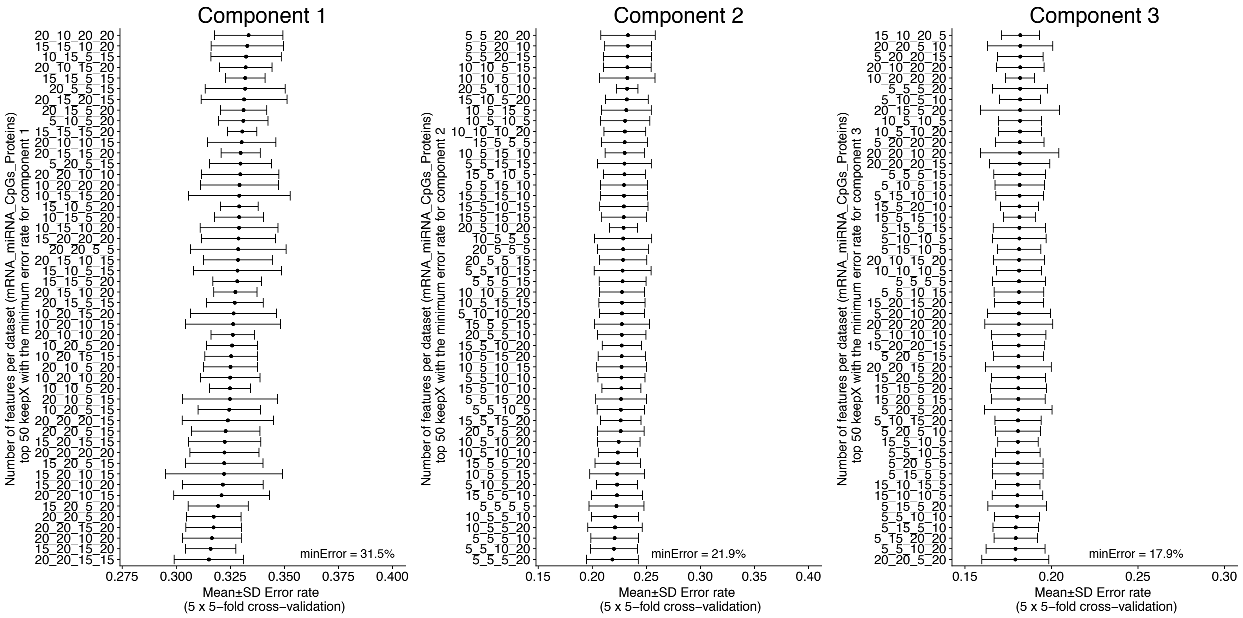

Supplementary Figure 8

**Breast cancer multi omics study: optimal multi-omics biomarker panel for PAM50 subtypes**. A grid was used to identify the optimal combination of variables select from each omics datasets. The following grid values was used for each omics dataset: mRNA = [5, 10, 15, 20], miRNA = [5, 10, 15, 20], CpGs = [5, 10, 15, 20], Proteins = [5, 10, 15, 20], across 3 components. The centroids distance measure was used to compute the error rate[26]. The optimal multi-omics panel consisted of 20 mRNAs, 20 miRNAs, 15 CpGs and 15 proteins on component 1, 5 mRNAs, 5 miRNAs, 5 CpGs and 20 proteins on component 2, and 20 mRNAs, 20 miRNAs, 5 CpGs and 20 proteins on component 3.

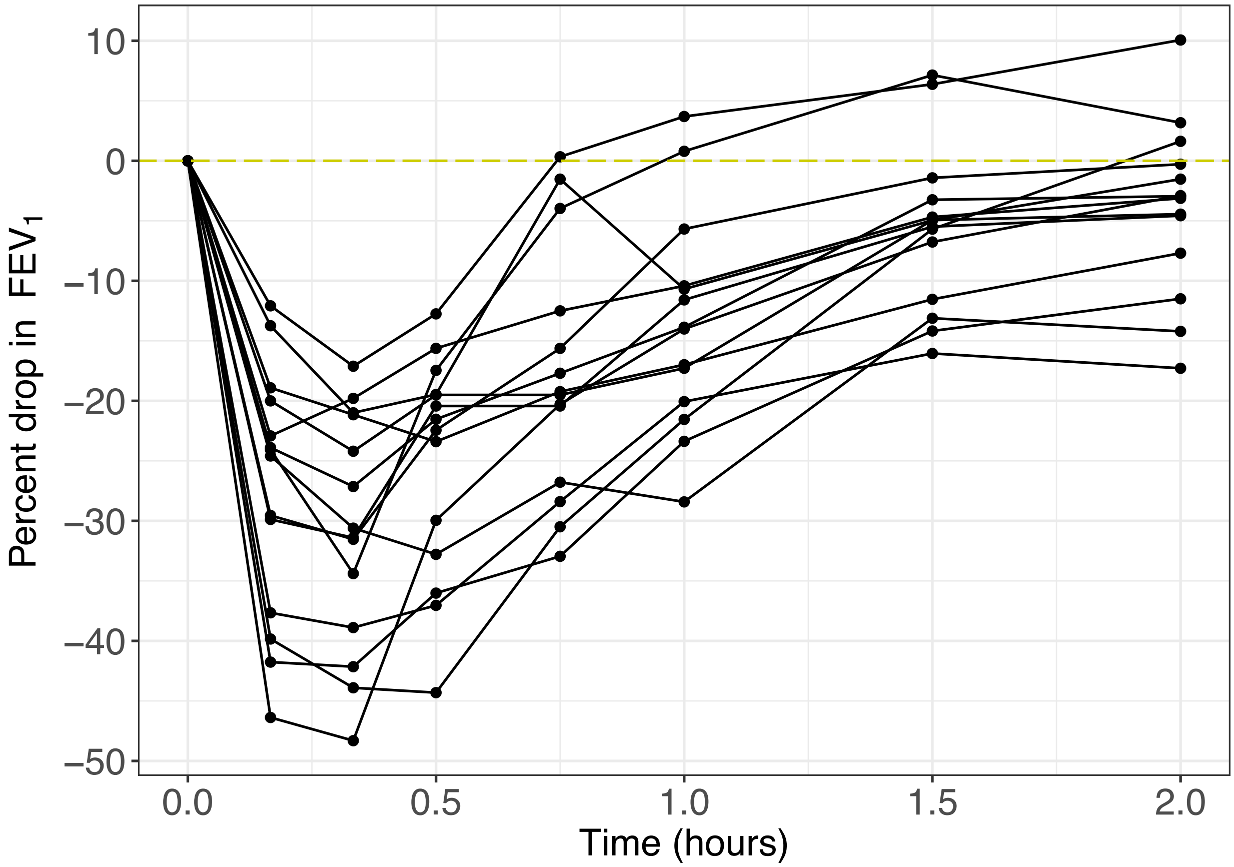

Supplementary Figure 9

**Asthma multi-omics study: decline in lung function after allergen inhalation challenge.**
Spirometry was used to measure the forced expiratory volume in one second of an exhale ($FEV_1$) prior to and at regularly interval after the allergen inhalation challenge.

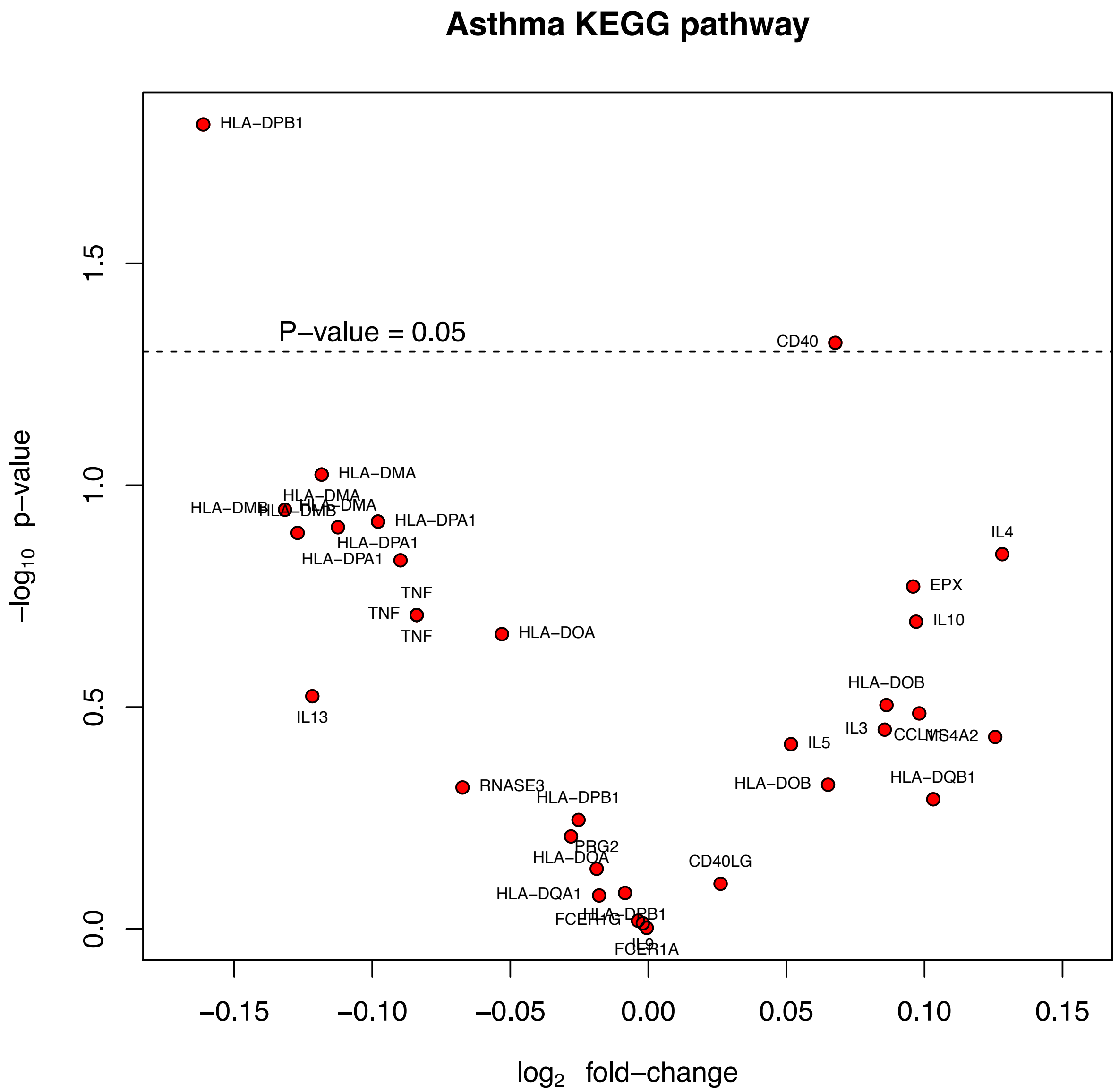

Supplementary Figure 10

**Asthma multi-omics study: volcano plot of genes in the Asthma KEGG pathway.** The volcano plot depicts the significance of each gene in the asthma pathways against its respective fold-change (change in expression from pre to-post challenge). The significance is based on a paired *t*-test. The volcano plot shows that with the exception of HLA-DPB1 and CD40 no other genes within the Asthma pathway were significant at the nominal p-value cut-off of 0.05. However, this pathway was selected by DIABLO as a strong predictor of allergen challenge. This modular-based analysis depicts the power of combining genes with small effect sizes which together contribute to a pathway that significantly changes in response to allergen inhalation challenge.

#### References

1. Huang, S., Chaudhary, K. & Garmire, L. X. More Is Better: Recent Progress in Multi-Omics Data Integration Methods. *Front. Genet.* **8,** (2017).
2. Akavia, U. D. *et al.* An Integrated Approach to Uncover Drivers of Cancer. *Cell* **143,** 1005–1017 (2010).
3. Zhu, J. *et al.* Stitching together multiple data dimensions reveals interacting metabolomic and transcriptomic networks that modulate cell regulation. *PLoS Biol.* **10,** e1001301 (2012).
4. Kim, D., Li, R., Dudek, S. M. & Ritchie, M. D. ATHENA: Identifying interactions between different levels of genomic data associated with cancer clinical outcomes using grammatical evolution neural network. *BioData Min.* **6,** 23 (2013).
5. Wang, W. *et al.* iBAG: integrative Bayesian analysis of high-dimensional multiplatform genomics data. *Bioinformatics* **29,** 149–159 (2013).
6. Lock, E. F., Hoadley, K. A., Marron, J. S. & Nobel, A. B. Joint and individual variation explained (JIVE) for integrated analysis of multiple data types. *Ann. Appl. Stat.* **7,** 523–542 (2013).
7. Zhang, S. *et al.* Discovery of multi-dimensional modules by integrative analysis of cancer genomic data. *Nucleic Acids Res.* **40,** 9379–9391 (2012).
8. Zhang, S., Li, Q., Liu, J. & Zhou, X. J. A novel computational framework for simultaneous integration of multiple types of genomic data to identify microRNA-gene regulatory modules. *Bioinformatics* **27,** i401–i409 (2011).
9. Argelaguet, R. *et al.* Multi-Omics factor analysis disentangles heterogeneity in blood cancer. *bioRxiv* 217554 (2017).
10. An Integrated Approach to Uncover Drivers of Cancer: Cell. Available at: http://www.cell.com/abstract/S0092-8674(10)01293-6. (Accessed: 12th February 2018)
11. Langfelder, P. & Horvath, S. WGCNA: an R package for weighted correlation network analysis. *BMC Bioinformatics* **9,** 559 (2008).
12. Wang, B. *et al.* Similarity network fusion for aggregating data types on a genomic scale. *Nat. Methods* **11,** 333–337 (2014).
13. Glass, K., Huttenhower, C., Quackenbush, J. & Yuan, G.-C. Passing messages between biological networks to refine predicted interactions. *PLoS ONE* **8,** e64832 (2013).
14. Lock, E. F. & Dunson, D. B. Bayesian consensus clustering. *Bioinformatics* **29,** 2610–2616 (2013).
15. Shen, H. & Huang, J. Sparse Principal Component Analysis via Regularized Low Rank Matrix Approximation. *J. Multivar. Anal.* **99,** 1015–1034 (2007).
16. Tenenhaus, A. *et al.* Variable selection for generalized canonical correlation analysis. *Biostatistics* **15,** 569–583 (2014).
17. González, I. *et al.* Highlighting relationships between heterogeneous biological data through graphical displays based on regularized canonical correlation analysis. *J. Biol. Syst.* **17,** 173–199 (2009).
18. van der Maaten, L. & Hinton, G. Visualizing Data using t-SNE. *J. Mach. Learn. Res.* **1,** 1–48 (2008).
19. Abdi, H., Williams, L. J. & Valentin, D. Multiple factor analysis: principal component analysis for multitable and multiblock data sets. *Wiley Interdiscip. Rev. Comput. Stat.* **5,** 149–179 (2013).

20. Zou, H. & Hastie, T. Regularization and variable selection via the elastic net. *J. R. Stat. Soc. Ser. B Stat. Methodol.* **67,** 301–320 (2005).
21. Lê Cao, K.-A., Boitard, S. & Besse, P. Sparse PLS discriminant analysis: biologically relevant feature selection and graphical displays for multiclass problems. *BMC Bioinformatics* **12,** 253 (2011).
22. Cun, Y. & Fröhlich, H. Network and data integration for biomarker signature discovery via network smoothed t-statistics. *PLoS ONE* **8,** e73074 (2013).
23. Sokolov, A., Carlin, D. E., Paull, E. O., Baertsch, R. & Stuart, J. M. Pathway-based genomics prediction using generalized elastic net. *PLoS Comput Biol* **12,** e1004790 (2016).
24. Breiman, L. Random forests. *Mach. Learn.* **45,** 5–32 (2001).
25. van de Wiel, M. A., Lien, T. G., Verlaat, W., van Wieringen, W. N. & Wilting, S. M. Better prediction by use of co-data: adaptive group-regularized ridge regression. *Stat. Med.* **35,** 368–381 (2016).
26. Rohart, F., Gautier, B., Singh, A. & Cao, K.-A. L. mixOmics: An R package for ‘omics feature selection and multiple data integration. *PLOS Comput. Biol.* **13,** e1005752 (2017).

### Supplementary Note

#### Real datasets

##### Benchmarking cancer datasets

All cancer (colon, glioblastoma, kidney and lung) datasets used for the benchmarking analyses were obtained from http://compbio.cs.toronto.edu/SNF/SNF/Software.html (Wang *et al.*[1]). For the mRNA datasets, all transcripts with the same gene symbol were averaged.

##### Breast cancer multi-omics study

*Datasets accession:* The level 3 TCGA data (version 2015_11_01) were retrieved from firebrowse.org hosted by the Broad Institute. The clinical data file (Merge_Clinical) was downloaded from the Primary tab of the BRCA Clinical Archives. The mRNA RSEM normalized dataset (illuminahiseq_rnaseqv2-RSEM_genes_normalized) was downloaded from the Primary tab of the BRCA mRNASeq Archives. The miRNA datasets (illuminahiseq_mirnaseq-miR_gene_expression and illuminaga_mirnaseq-miR_gene_expression) were downloaded from the Primary tab of the BRCA miRSeq Archives. The reverse phase protein array dataset (mda_rppa_core-protein_normalization) was downloaded from the Primary tab of the BRCA RPPA Archives. The beta values for the methylation datasets (humanmethylation27-within_bioassay_data_set_function and humanmethylation450-within_bioassay_data_set_function MD5) were downloaded from the Primary tab of the BRCA Methylation Archives.

*Data processing:* Clinical data were present for 1,098 subjects for 3,703 variables. 29 unannotated transcripts were removed from the mRNA dataset composed resulting in 20,502 genes x 1212 samples. Two transcripts corresponded to *SLC35E2*, therefore one of the transcripts was re-labelled *SLC35E2.rep*. The miRNA datasets (1,046 miRNA x 1190 samples) was derived using two different Illumina technologies, the Illumina Genome Analyzer (341 samples) and the Illumina HiSeq (849 samples). The read counts instead of the reads_per_million_miRNA_mapped were used. The proteomics dataset obtained using a reverse phase protein array consisted of 142 proteins for 410 samples. The methylation data was derived from two different platforms, the Illumina Methylation 27 (27,578 CpG probes x 343 subjects) and the Illumina 450K (485,577 CpG probes x 885 subjects). There were 25,978 CpG probes in common between the platforms. The PAM50 labels for 1,182 samples were obtained from the TCGA staff. All datasets were restricted to samples coming from the primary solid tumor (sample type code 01) and to the first vial (vial code A).

*Normalization and pre-filtering:* The count data for the mRNA dataset was normalized to log2-counts per million (logCPM), similar to limma voom [2]:

$$X_{norm} = \log_2\left(\frac{\left(X_{counts} + 0.5\right)^T}{\left(lib.size + 1\right) * 10^6}\right)$$

After library size normalization, genes with counts less than 0 were removed. The PAM50 genes were also removed from the mRNA dataset prior to analyses. Similarly, the miRNA count data was normalized to logCPM and miRNA transcripts with counts less than 0 were also removed.

#### Asthma multi-omics study

*Datasets accession:* Paired blood samples were obtained from 14 asthmatic individuals undergoing allergen inhalation challenge as previously described [3]. Cell counts were obtained from a hematolyzer (percentage of Neutrophils, Lymphocytes, Monocytes, Eosinophils and Basophils) and DNA methylation analysis (percentage of T regulatory cells, T cells, B cells and Th17 cells). Gene expression profiling was performed using Affymetrix Human Gene 1.0 ST (GSE40240). Metabolite profiling was performed by Metabolon Inc. (Durham, North Carolina, USA). All asthma data have been published as part of previous studies [4,5].

*Normalization:* Microarray data was normalized using Robust MultiArray Average (RMA), consisting of background correction, quantile normalization and probe summarization using median polish. Preprocessing of mass spectrometry data including data extraction, peak-identification and data preprocessing for quality control and compound identification was performed by Metabolon Inc. (Durham, North Carolina, USA).

#### Simulated datasets

##### Generating multi-omics data

Three datasets were simulated each with 200 observations (n) and 260 variables (p). The 200 observations were split equally over two groups (G1 and G2), whereas the 260 variables were generated by varying the degree of correlation and fold-change (δ) between G1 and G2: 30 correlated-discriminatory (corDis) variables, 30 uncorrelated-discriminatory (unCorDis) variables, 100 correlated-nondiscriminatory (corNonDis) variables, and 100 uncorrelated-nondiscriminatory (unCorNonDis) variables. The resulting dataset was of the form:

$$\boldsymbol{X}_j = [\boldsymbol{X}_j^{corDis} \quad \boldsymbol{X}_j^{unCorDis} \quad \boldsymbol{X}_j^{corNonDis} \quad \boldsymbol{X}_j^{unCorNonDis}] + \boldsymbol{E}_j, \quad j = 1, \ldots, 3.$$

The discriminatory variables (corDis and unCorDis) were generated using the following model:

$$\boldsymbol{X}_j = \boldsymbol{u}_j \boldsymbol{w}_j^t, \text{ where } ||\boldsymbol{w}|| = 1 \quad j = 1, \ldots, 3,$$

where the loadings, $\mathbf{w}_1$, $\mathbf{w}_2$, and $\mathbf{w}_3$ were 30-vectors, and the elements were drawn from a uniform distribution in the interval of [-0.3, 0.2] U [0.2, 0.3]. For G1, the outer components $\mathbf{u}_1$, $\mathbf{u}_2$, $\mathbf{u}_3$ were 3-vectors drawn from a multivariate normal distribution with a mean value of -δ/2 and a mean value of δ/2 for G2. For corDis variables, cor($\mathbf{u}_1$, $\mathbf{u}_2$) = 1, cor($\mathbf{u}_1$, $\mathbf{u}_3$) = 1, cor($\mathbf{u}_2$, $\mathbf{u}_3$) = 1, whereas for unCorDis variables, cor($\mathbf{u}_1$, $\mathbf{u}_2$) = 0, cor($\mathbf{u}_1$, $\mathbf{u}_3$) = 0, cor($\mathbf{u}_2$, $\mathbf{u}_3$) = 0.

The nondiscriminatory variables (corNonDis and unCorNonDis) were generated by drawing 100-vectors each with 200 elements, from a multivariate normal distribution with a mean of 0. For corNonDis variables, cor($\mathbf{u}_1$, $\mathbf{u}_2$) = 1, cor($\mathbf{u}_1$, $\mathbf{u}_3$) = 1, cor($\mathbf{u}_2$, $\mathbf{u}_3$) = 1, whereas for unCorNonDis variables, cor($\mathbf{u}_1$, $\mathbf{u}_2$) = 0, cor($\mathbf{u}_1$, $\mathbf{u}_3$) = 0, cor($\mathbf{u}_2$, $\mathbf{u}_3$) = 0.

$\boldsymbol{E}_j$ is a 200 x 260 residual matrix where each element is drawn from a normal distribution with zero mean and variance according to the grid [0.1, 0.2, 0.6, 1]. The following grid of values were used for the fold-change: [0.1, 0.5, 1, 2].

#### Simulation analysis

Using fold-change values of [0.5, 1, 2] and noise values of [0.2, 0.5, 1, 2], 16 (4x4) sets of three datasets were generated, and DIABLO was applied, either with the full or null design (DIABLO_full and DIABLO_null). The full design, connects all blocks in the design matrix (C), such that $c_{ij}$=1, i=1,2,3 and j=1,2,3, whereas the null design does not connect any datasets in the design matrix (C), such that cij=0, i=1,2,3 and j=1,2,3. One component was retained in the DIABLO model, selecting 30 variables from each dataset for a total of 90 variables (across all datasets). In addition, other integrative schemes such as concatenation and ensemble-based classifiers were also tested using the sPLSDA classifier. For the concatenation-based scheme, all datasets were concatenated into one matrix containing 3x260=880 variables and sPLSDA was applied, retaining 1 component and 90 variables. For the ensemble-based scheme, a sPLSDA classifier was applied to each dataset separately retaining one component and 30 variables per dataset. The consensus predictions were determined using a majority vote scheme. A 10-fold cross-validation averaged over 50 simulations was used to evaluate the performance of each method/scheme and the number of each type of variable selected in each model was recorded.

#### Description of methods used for the benchmarking experiments

For the purposes of this study, only component-based methods that integrated multiple datasets and perform variable selection were considered. Since tuning the number of variables to retain in each model would result in biomarker panels with different numbers of variables, for the purposes of this study all variables were retained in each model. The features were instead ranked based on their absolute value of their loadings (importance) and 60 variables were selected from each omic type, resulting in multi-omic biomarker panels with 180 variables (60 mRNAs, 60 miRNAs and 60 CpGs). Equal numbers of variables allowed for a fair comparison in the gene set enrichment analysis.

| | **Parameter settings** |
|---|---|
| **Supervised** | |
| DIABLO_null | `ncomp = 2` (# of components)<br>`keepX` = all variables were retained from each omics dataset<br>$design = \begin{bmatrix} 0 & 0 & 0 \\ 0 & 0 & 0 \\ 0 & 0 & 0 \end{bmatrix}$<br><br>default parameters were used for the other arguments:<br>`scheme="horst",`<br>`mode="regression",`<br>`scale = TRUE,`<br>`init = "svd",`<br>`tol = 1e-06,`<br>`max.iter = 100` |
| DIABLO_full | `ncomp = 2` (# of components)<br>`keepX` = all variables were retained from each omics dataset |

| | |
|---|---|
| | $design = \begin{bmatrix} 0 & 1 & 1 \\ 1 & 0 & 1 \\ 1 & 1 & 0 \end{bmatrix}$<br><br>default parameters were used for the other arguments:<br>`scheme="horst",`<br>`mode="regression",`<br>`scale = TRUE,`<br>`init = "svd",`<br>`tol = 1e-06,`<br>`max.iter = 100` |
| Concatenation-sPLSDA | `ncomp = 2` (# of components)<br>`keepX` = all variables were retained from each omics dataset<br><br>default parameters were used for the other arguments:<br>`mode = "regression"`<br>`scale = TRUE,`<br>`tol = 1e-06,`<br>`max.iter = 100` |
| Ensemble_sPLSDA | `ncomp = 2` (# of components)<br>`keepX` = all variables were retained from each omics dataset<br><br>default parameters were used for the other arguments:<br>`mode = "regression"`<br>`scale = TRUE,`<br>`tol = 1e-06,`<br>`max.iter = 100` |
| **Unsupervised** | |
| sGCCA[6] | `ncomp = 2` (# of components)<br>`keepX` = all variables were retained from each omics dataset<br>$design = \begin{bmatrix} 0 & 1 & 1 \\ 1 & 0 & 1 \\ 1 & 1 & 0 \end{bmatrix}$<br><br>default parameters were used for the other arguments:<br>`scheme = "horst",`<br>`mode="canonical",`<br>`scale = TRUE,`<br>`init = "svd.single",`<br>`tol = .Machine$double.eps,`<br>`max.iter=1000,` |
| JIVE*[7] | default parameter settings from the jive() from the r.jive R-package were used:<br>1. `scale = TRUE, center = TRUE` |

| | |
|---|---|
| | 2. `method = “perm”`<br><br>sPCA parameters:<br>`ncomp = 2` (# of components)<br>`keepX = rep(ncol(X),ncomp)`(all variables were retained from each omics dataset<br><br>default parameters were used for the other arguments:<br>`center = TRUE`<br>`scale = TRUE,`<br>`max.iter = 500,`<br>`tol = 1e-06` |
| MOFA[8] | `factors=2` (# of components)<br><br>default parameter settings recommended by MOFA were used:<br>1. `likelihoods=( gaussian gaussian gaussian )`<br>2. Convergence criterion (`tolerance=0.01, nostop=0`)<br>3. Training components (`startDrop=1` # initial iteration to start shutting down factors, `freqDrop=1` # frequency of checking for shutting down factors, `dropR2=0.00` # threshold on fraction of variance explained)<br>4. hyperparameters for the feature-wise spike-and-slab sparsity prior [`learnTheta=( 1 1 1 )` # 1 means that sparsity is active whereas 0 means the sparsity is inactivated; each element of the vector corresponds to a view, `initTheta=( 1 1 1 )` # initial value of sparsity levels (1 corresponds to a dense model, 0.5 corresponds to factors ); each element of the vector corresponds to a view, `startSparsity=250` # initial iteration to activate the spike and slab, we recommend this to be significantly larger than 1]<br>Intercept was set to TRUE (`learnIntercept=1`) |

*since the variable selection functionality has not been added to JIVE R-function, sparse Principal Component Analysis (sPCA) from the mixOmics R-package was applied to the joint variation matrix obtained after applied JIVE to the multi-omics cancer datasets.

#### References

1. Wang, B. *et al.* Similarity network fusion for aggregating data types on a genomic scale. *Nat. Methods* **11,** 333–337 (2014).
2. Law, C. W., Chen, Y., Shi, W. & Smyth, G. K. Voom: precision weights unlock linear model analysis tools for RNA-seq read counts. *Genome Biol* **15,** R29 (2014).
3. Singh, A. *et al.* Plasma proteomics can discriminate isolated early from dual responses in asthmatic individuals undergoing an allergen inhalation challenge. *PROTEOMICS - Clin. Appl.* **6,** 476–485 (2012).
4. Singh, A. *et al.* Gene-metabolite expression in blood can discriminate allergen-induced isolated early from dual asthmatic responses. *PLoS ONE* **8,** e67907 (2013).
5. Singh, A. *et al.* Th17/Treg ratio derived using DNA methylation analysis is associated with the late phase asthmatic response. *Allergy Asthma Clin. Immunol.* **10,** 32 (2014).
6. Tenenhaus, A. *et al.* Variable selection for generalized canonical correlation analysis. *Biostatistics* **15,** 569–583 (2014).
7. Lock, E. F., Hoadley, K. A., Marron, J. S. & Nobel, A. B. Joint and individual variation explained (JIVE) for integrated analysis of multiple data types. *Ann. Appl. Stat.* **7,** 523–542 (2013).
8. Argelaguet, R. *et al.* Multi-Omics factor analysis disentangles heterogeneity in blood cancer. *bioRxiv* 217554 (2017).
